# Endothelial β3-Adrenergic Receptor activation prevents pulmonary hypertension

**DOI:** 10.64898/2026.07.14.738203

**Authors:** Susana F. Rocha, Laura de la Bastida-Casero, Monika Spaczyńska-Kwiatkowska, Álvaro Macías, Yolanda Sierra-Palomares, Mónica Gomez, Anabel Díaz-Guerra, María Villaba-Orero, Víctor I. Peinado, Ana García-Álvarez, Joan Albert Barberá, Valentín Fuster, Borja Ibañez, Eduardo Oliver

**Affiliations:** Centro Nacional de Investigaciones Cardiovasculares (CNIC), Madrid, Spain; Centro de Investigaciones Biológicas Margarita Salas (CIB-CSIC), Madrid Spain; Department of Physiotherapy, Faculty of Medicine, Health and Sports, Universidad Europea de Madrid, Madrid, Spain; Centro Nacional de Investigaciones Oncológicas, Madrid, Spain; Centro de Investigaciones Biomédicas en Red de Enfermedades Cardiovasculares (CIBERCV), Madrid, Spain; Departamento de Medicina y Cirugía Animal, Facultad de Veterinaria, Universidad Complutense de Madrid, Madrid, Spain; Instituto de Investigaciones Biomédicas de Barcelona (IIBB-CSIC-IDIBAPS), Barcelona, Spain; Hospital Clínic de Barcelona, Institut d’Investigacions Biomèdiques August Pi i Sunyer (IDIBAPS), University of Barcelona, Barcelona, Spain; Centro de Investigaciones Biomédicas en Red de Enfermedades Respiratorias (CIBERES), Madrid, Spain; Icahn School of Medicine at Mount Sinai, NY, USA; IIS-Fundación Jiménez Díaz University Hospital, Madrid, Spain; Max-Planck Institute for Molecular Biomedicine, Münster, Germany

**Keywords:** beta3-adrenergic receptor, pre-capillary pulmonary hypertension, endothelial dysfunction, mitochondrial fitness

## Abstract

**Background:** Pulmonary hypertension (PH) is a progressive vascular disease characterized by endothelial dysfunction, vascular remodeling and increased pulmonary vascular resistance. The β3-adrenergic receptor (β3-AR) has been implicated in cardiovascular regulation and cardioprotective mechanisms; however, its role in pulmonary vascular disease remains poorly understood. We investigated whether activation of β3-AR protects pulmonary endothelial function and prevents the development of pre-capillary PH.

**Methods:** β3-AR expression was evaluated in pulmonary endothelium from patients with Chronic Obstructive Pulmonary Disease (COPD) and in murine models of hypoxia-induced PH. Genetic mouse models including β3-AR knockout (KO) and conditional β3-AR overexpression in endothelial cells (EC) or in smooth muscle cells (SMC), were used to determine cell-specific roles. Pharmacological activation of β3-AR was achieved using the selective β3-agonist mirabegron in hypoxia-induced PH mice and monocrotaline-induced PH rats. Pulmonary vascular reactivity and vasodilatory responses to β3-AR stimulation were evaluated by wire myography in isolated pulmonary arteries. Mechanistic studies were performed in human pulmonary artery endothelial cells (HPAEC) under hypoxic conditions, in human pulmonary arterial smooth muscle cells (HPASMC) and in endothelial nitric oxide synthase (NOS3) KO mice.

**Results:** β3-AR was upregulated in pulmonary endothelium of COPD patients and mice exposed to chronic hypoxia. Genetic deletion of β3-AR aggravated PH, whereas endothelial-specific overexpression attenuated the disease phenotype, reducing right ventricular systolic pressure (RVSP), vascular remodeling and right ventricular (RV) hypertrophy. Activation of β3-AR with mirabegron improved pulmonary hemodynamics, reduced vascular remodeling and preserved RV function. β3-AR activation promoted endothelial nitric oxide synthase (eNOS)-dependent NO production, indirectly inhibiting SMC proliferation. Additionally, β3-AR activation improved mitochondrial fitness in endothelial cells by increasing uncoupling protein 2 (UCP2) expression, reducing reactive oxygen species (ROS) generation and preventing mitochondrial fragmentation.

**Conclusions:** These findings identify endothelial β3-AR as a previously unrecognized regulator of pulmonary vascular homeostasis and provide a strong translational rationale for targeting the β3-adrenergic pathway in PH. Given that mirabegron is already approved for clinical use, our results support its repurposing as a therapeutic strategy for pre-capillary forms of PH.

## INTRODUCTION

Pulmonary hypertension (PH) is a group of aggressive multifactorial diseases that are characterized by a chronic elevation of the mean pulmonary arterial pressure (mPAP) above 20 mmHg at rest. The etiology of PH is very heterogeneous and varies from idiopathic and hereditary to environmental factors, such as hypoxia or infectious agents, in addition to secondary causes related to cardiac and lung pathologies.^1–4^ The most recent guidelines classify PH into 5 groups depending on its origin, as well as different pathophysiological, clinical and therapeutic considerations: (1) Pulmonary Arterial Hypertension – PAH; (2) PH associated with left-sided heart disease – LHD-PH; (3) PH associated with chronic lung disease or hypoxia – CLD-PH; (4) Chronic Thromboembolic PH – CTEPH; and (5) PH with unclear or multifactorial origin. Hemodynamically, PH is further classified as pre-capillary, isolated post-capillary (Ipc-PH) and, as the diseases evolves, combined pre- and post-capillary (Cpc-PH).^4^ Pre-capillary PH, which encompasses PAH, CLD-PH or CETPH, is defined by mPAP >20 mmHg at rest, pulmonary capillary wedge pressure (PCWP) ≤15 mmHg and a pulmonary vascular resistance (PVR) >2 Woods units.^4^

This characteristic chronic increase in mPAP and PVR results from functional and structural changes in small pulmonary arterioles, initiated by endothelial damage and dysfunction. This leads to an imbalance in vasoactive mediators and growth factors.^5^ This dysregulation impairs vasodilation, increases vascular tone, and promotes oxidative stress, inflammation, fibrosis, coagulation, and vascular cell proliferation.^5,6^ Mitochondrial dysfunction and an imbalance between apoptosis and proliferation further contribute to cellular stress and metabolic reprogramming.^5^ Together, these processes drive pulmonary vascular thickening, plexiform lesion formation, vascular obliteration, and increased pulmonary pressure.^7,8^ The resulting chronic pressure overload induces progressive right ventricular (RV) remodeling, initially as compensatory hypertrophy and ultimately leading to RV dysfunction, heart failure, and premature death.^9^ Although the development of specific treatments has notably improved the prognosis of PH, none of them has proved to be sufficiently effective to tackle both cellular and structural problems of the vasculature.^7,8,10^ More recently, drugs targeting vascular remodeling, such as Sotatercept, have demonstrated significant clinical benefit in PAH patients,^11^ however, other forms such as CLD-PH or CTEPH still lack of effective therapies.^12,13^

Beta-adrenergic receptors (β-AR) are members of the G-proteins-coupled receptors family (GPCR), widely distributed in the organism, that are involved in the cellular response and of special interest for the cardiovascular system. Among the three β-AR subtypes, β1-AR and β2-AR subtypes were for long time thought to be the only ones existing in the heart and airways respectively.^14^ Later, β3-AR that was first discovered in adipocytes and to be involved in metabolism was also found in the heart and vessels in different conditions.^15^ Vascular β3-AR was found both in smooth muscle cells (SMCs) and endothelial cells (ECs), depending on the territory, where controls vasorelaxation either via cAMP/PKA or activation of endothelial nitric oxide synthase (eNOS).^16–18^ Although β3-AR levels in physiological conditions are low in the cardiovascular system, they undergo up-regulation in certain pathological conditions having been suggested to be a protective mechanism.^19–23^ In this scenario, stimulation of the β3 adrenergic pathway has been proposed as a potential treatment for vascular and cardiac pathologies such as HF,^18,19^ aortic stenosis^24^, or myocardial infarct among others.^23,25^ However, despite the use of β3-agonist has been previously suggested in certain models of PH,^21,26^ its cellular and molecular mechanism has not been elucidated, which is necessary to define patients that would better benefit from this therapy.

In this work we used human lung samples and different transgenic models to demonstrate that β3-AR has an essential and multifactorial role in PH, by regulating EC metabolism and function, controlling pulmonary vascular remodeling and contractility, and ultimately improving RV hemodynamics and attenuating RV dysfunction. All this is done following modulation of mitochondrial ultrastructure and fitness, NO and ROS production, both via eNOS and the mitochondrial uncoupling protein 2 (UCP2) pathways. We also demonstrate that the use of the β3-adrenergic agonist mirabegron, a Food and Drug Administration (FDA) approved therapy to treat overactive bladder, is protective in two different rodent models of pre-capillary PH. Our results lead us to consider β3-AR as an excellent therapeutic target, and the β3-adrenergic agonist mirabegron as a potential therapeutic strategy to treat pre-capillary forms of PH such as PAH or CLD-PH among others in which regulation of endothelial function plays an essential role.

## METHODS

Supplemental Methods are provided in the Supplemental Material.

### Animal models

All animal procedures complied with European and Spanish regulations, were approved by the local ethics committees (CSIC and Universidad Autónoma de Madrid-CNIC) and the Animal Protection Area of the Comunidad Autónoma de Madrid (PROEX 112.5/20 and PROEX 195.2/22), and conformed to ARRIVE guidelines 2.0. The role of β3-AR signaling in PH was investigated using global β3-AR knockout mice (AdrB3^KO^) and genetic rescue models with EC- or SMC-specific restoration of the β3-AR expression (AdrB3^EC^ and AdrB3^SMC^) compared to litter mate control mice. Additional experiments were performed in eNOS KO (NOS3^KO^) and wild-type (WT) mice. PH was also induced in C57Bl/6 WT mice by chronic hypoxia exposure (2 weeks at 10% O_2_) and in Sprague Dawley rats by monocrotaline (MCT) administration. The selective β3-AR agonist mirabegron was administered (2 and 10 mg/kg/day) in drinking water either during hypoxia exposure in mice or using preventive and therapeutic treatment protocols in monocrotaline-challenged rats. Pulmonary hemodynamics, right ventricular remodeling and cardiac function were assessed by invasive pressure measurements, Fulton’s index and transthoracic echocardiography. Pulmonary vascular remodeling was quantified in lung sections by histology and immunostaining, and pulmonary artery function was evaluated by wire myography. All animal experiments were conducted by experienced researchers blinded to experimental groups.

### Human samples

Human lung samples were obtained from lung transplant recipients with chronic obstructive pulmonary disease (COPD) with or without PH.^27^ Control samples were obtained from donors undergoing surgical lung resection for lung cancer as part of a study of histology in smokers.^28^ Lung tissue samples were collected prospectively and processed after surgery. Studies were approved by the Ethics Committee of Hospital Clinic, Barcelona, Spain, and all patients provided a written informed consent.

### Cell culture and molecular biology

Mechanistic studies were performed in human pulmonary arterial endothelial (HPAEC) and smooth muscle cells (HPASMC) exposed to normoxia (21% O_2_) or hypoxia (2% O_2_ for 2 and 24h) and treated with mirabegron, with or without inhibitors of NOS or UCP2. Endothelial–SMC crosstalk, SMC proliferation, oxidative stress, mitochondrial content, mitochondrial respiration and molecular signaling were analyzed using conditioned-medium experiments, EdU assays, fluorescence probes, immunofluorescence, Seahorse extracellular flux analysis, western blotting, flow cytometry, quantitative PCR and ELISA.

### Statistics

Data are shown as mean ± SEM. Statistical analyses were performed using GraphPad Prism. Two-group comparisons were analyzed by two-tailed Student’s t test, and multiple-group comparisons by ANOVA followed by appropriate post hoc tests, as specified in the figure legends. P<0.05 was considered statistically significant.

## RESULTS

### β3-AR is overexpressed in human and mice pulmonary arterioles affected by chronic hypoxic conditions

To investigate the potential implication of β3-AR in hypoxia-related diseases, we measured its expression in lung tissue from patients with COPD, a condition characterized by alveolar hypoxia.^29^ We found that patients with COPD, which frequently develop group 3 PH,^30^ present increased β3-AR protein levels in the lung endothelium that correlate with a worse prognosis measured by the BODE index (Body-mass index, airflow Obstruction, Dyspnea, and Exercise), a predictor of the number and severity of acute COPD exacerbations (Fig. 1A-C). Following this result, we wanted to explore whether β3-AR plays a role in pre-capillary forms of PH. For this, we first assessed β3-AR activity in pulmonary arteries from mice subjected to chronic hypoxia (Fig. 1D). This was achieved *ex vivo,* by stimulating phenylephrine pre-contracted normoxic or hypoxic pulmonary arteries in a wired myograph with different concentrations of the selective β3-agonist, mirabegron. Whereas normoxic WT arteries failed to vasodilate in response to mirabegron-mediated β3-AR stimulation, mirabegron induced a potent vasodilator effect in vessels from PH mice (Fig. 1E, compare black and red fitted lines). To confirm that this mirabegron-mediated vasodilation was specific, we also performed experiments using pulmonary arteries from AdrB3^KO^. Concentration-response curves showed that the vasodilator effect of mirabegron was decreased by 75% in AdrB3^KO^ compare to hypoxic WT mice, as evidenced by a shift of the concentration-response curve (Fig. 1E, green fitted lines). These results show a clear upregulation of β3-AR upon hypoxia exposure and support the β3-AR-specific vasodilator effect of mirabegron in hypoxic pulmonary arteries.

**Figure 1.**
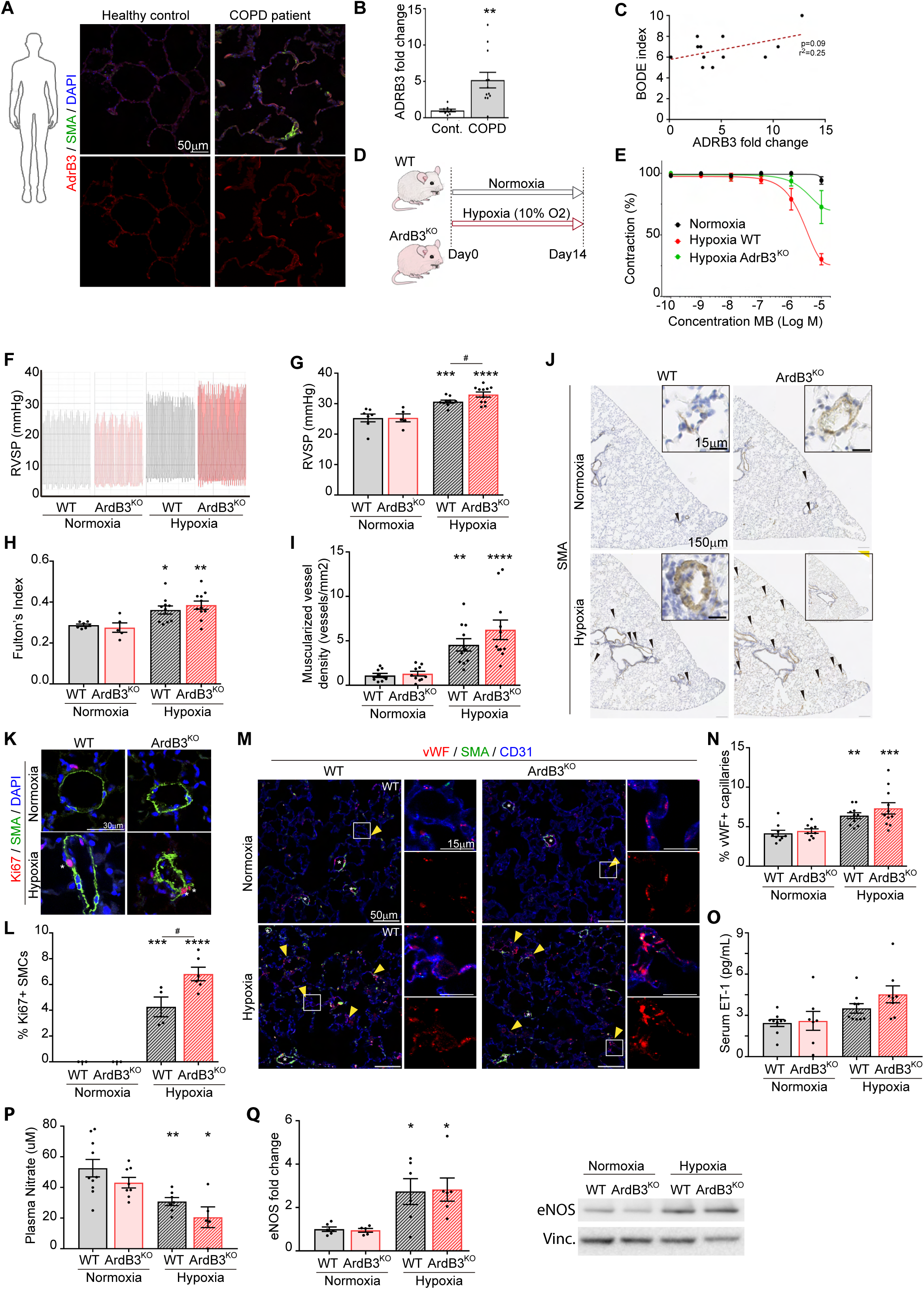
β3-AR expression is increased in pulmonary vessels of COPD patients and its absence aggravates hypoxia-induced pulmonary hypertension and endothelial dysfunction in mice. **(A)** Representative confocal images of β3-AR (red) and SMA (green) immunostaining in human lung samples from healthy and COPD patients. **(B)** Quantification of fold change in percentage of lung area with β3-AR (ADRB3) immunostaining. **(C)** Correlation between fold change in β3-AR protein expression in COPD patients and BODE index (the higher the BODE index, the poorer prognosis). **(D)** Hypoxia-induced pulmonary hypertension experimental setup. **(E)** Relaxation curves of phenylephrine pre-contracted normoxic (black line) or hypoxic pulmonary arteries of WT (red line) or AdrB3^KO^ mice (green line) in a wired myograph after different concentrations of the selective β3-AR-agonist, mirabegron. **(F, G)** Representative recording profiles and quantification of RVSP. (H) Fulton’s index reflecting RV hypertrophy. **(I)** Representative images of mouse lung sections immunostained for SMA (smooth muscle actin). Arrowheads indicate fully muscularized arterioles with less than 30um diameter. Inserts depict representative arterioles. **(J)** Quantification of density of fully muscularized arterioles in lung sections. **(K)** Representative confocal images of Ki67 (red) and SMA (green) immunostaining in mouse lung sections. Asterisks indicate proliferative Ki67^+^ SMCs. **(L)** Quantification of proliferating lung SMCs. **(M)** Representative confocal images of vWF (red), SMA (green) and CD31 (blue) immunostaining in mouse lung sections. Asterisks indicate arterioles, where vWF expression is observed in both normoxic and hypoxic conditions and yellow arrowheads indicate ectopic expression of vWF in capillaries. Insets of capillary regions are shown to better appreciate ectopic vWF expression. **(N)** Quantification of percentage of capillary bed with vWF expression. **(O)** Quantification of circulating ET-1 levels. (P) Quantification of circulating nitrate as a readout of nitric oxide. **(Q)** Quantification of eNOS protein levels in lungs and representative image of western blot. All data are represented as mean ± s.e.m. Each dot in charts represents an individual patient/mouse. Two-tailed, unpaired t-test and two-way ANOVA with Sidak’s multiple comparisons test for remaining charts. Asterisks indicate p-values between normoxic and hypoxic mice of the same genotype and hashtags indicate p-values between genotypes. *p<0.05, **p<0.01, ***p<0.005, ****p<0.001, # p <0.05.

### Absence of β3-AR exacerbates hypoxia-induced PH

In order to further evaluate the role of β3-AR in the development of PH, we assessed hemodynamic and morphological variables such as RVSP, RV hypertrophy (Fulton’s index) and pulmonary vascular remodeling of WT and AdrB3^KO^ mice after 2 weeks of hypoxic exposure (Fig. 1F-K; Sup. Fig. 1A-C). In normoxic conditions, WT and AdrB3^KO^ mice presented no differences in RVSP (25.33 ± 1.29 mmHg and 25.34 ± 1.30 mmHg, respectively), however upon chronic hypoxia and the induction of PH, AdrB3^KO^ mice presented a worsen phenotype with significantly augmented RVSP compared to WT mice (33.01 ± 0.50 mmHg in AdrB3^KO^ mice and 30.71 ± 0.47 mmHg in control mice; p=0.032; Fig. 1F-G). A similar trend was observed in the Fulton’s index (hypoxic WT mice with 0.361 ± 0.020 vs hypoxic AdrB3^KO^ with 0.385 ± 0.020; Fig 1H). Pulmonary vascular remodeling was assessed both by quantifying the vessel density of fully muscularized vessels (SMA+ and with diameter <30μm) and percentage of proliferative SMCs (Fig. 1I-J). AdrB3^KO^ mice tend to have more remodeled vessels than WT mice in hypoxia (hypoxic WT with 4.55 ± 0.69 vessels/mm^2^ vs hypoxic AdrB3^KO^ mice with 6.25 ± 1.10 vessels/mm^2^; Fig. 1 I-J). Accordingly, we observed that pulmonary arterioles from hypoxic AdrB3^KO^ mice present a significantly higher percentage of proliferating SMCs (Ki67+, SMA+ cells) than those from hypoxic WT mice (hypoxic WT with 4.27 ± 0.77% vs hypoxic AdrB3^KO^ mice with 6.81 ± 0.53%; p=0.021; Fig. K-L), while proliferative SMCs are absent in normoxic arterioles.

Given that AdrB3^KO^ mice present a more severe PH phenotype with increased vascular remodeling when compared to WT mice, we questioned if endothelial dysfunction was also exacerbated in the absence of β3-AR. Relevant markers of endothelial damage/dysfunction, such as, ectopic vWF expression in capillaries (Fig. 1M-N) and circulating endothelin (ET-1; Fig. 1O), tend to be increased in hypoxic AdrB3^KO^ mice when compared to hypoxic WT controls. Also, in accordance with increased endothelial damage in the absence of β3-AR, hypoxic AdrB3^KO^ mice also present decreased NO production, despite eNOS protein levels being as elevated as control hypoxic mice (Fig. 1P-Q). These findings suggest a functional impairment of the eNOS-NO axis rather than a reduction in eNOS protein expression. CD31, an endothelial cell adherence junction protein important for its integrity, is also decreased in hypoxic AdrB3^KO^ mice (Sup. Fig. 1D). In order to discard that changes were due to a reduction in EC content, we performed FACS-analysis of lung tissue. Results show that EC content is not reduced but nearly duplicates in both control and AdrB3^KO^ hypoxic mice (Sup. Fig. 1E-F). Altogether, these results indicate that β3-AR has a prominent role in protecting the endothelium from hypoxia-induced damage.

### Restoration of endothelial β3-AR ameliorates the hypoxia-induced PH phenotype in KO mice

To explore whether restoration of endothelial β3-AR would be sufficient to prevent endothelial damage and PH, we interbred AdrB3^KO^ and R26^LSL-Β3-AR-IRES-GFP^Tie2^Cre^ mice^24,25^ that conditionally overexpress human β3-AR in the endothelial compartment (Fig. 2A-B; Sup. Fig.2A). Using three different techniques—confocal immunofluorescence imaging, flow cytometric assessment of transgene expression, and gene expression analysis of FACS-sorted endothelial cells—we confirmed successful human β3-AR expression in pulmonary ECs (Fig. 2A, C, D), and that levels of the resulting offspring (hereafter named AdrB3^EC^), are similar to that of lung ECs of WT in normoxia (Fig. 2A, D). Restoration of endothelial β3-AR expression was sufficient to significantly attenuate RVSP in AdrB3^EC^ when compared to AdrB3^KO^ littermates subjected to chronic hypoxia (hypoxic AdrB3^KO^ with 28.12 ± 0.53 mmHg vs hypoxic AdrB3^EC^ mice with 23.77 ± 0.56 mmHg; p<0.0001) and also the Fulton’s index (hypoxic AdrB3^KO^ mice with 0.422 ± 0.018 vs hypoxic AdrB3^EC^ with 0.356 ± 0.008; p<0.01) (Fig. E-G; Sup. Fig. 2B-D). In addition to the ameliorated hemodynamic variables, restoration of endothelial β3-AR expression also led to a dramatic decrease in SMC proliferation (hypoxic AdrB3^KO^ with 9.28 ± 2.06 % vs hypoxic AdrB3^EC^ mice with 1.26 ± 0.90 % of Ki67+ SMCs; p<0.005; Fig 2H-I), consistent with presence of thinner vessels. No differences were observed in the frequency of remodelled vessels (Fig. 2J-K). Additionally, we also generated mice with restored human β3-AR expression in SMCs (Sup. Fig. 3A), which is a major component of arteries/arterioles (hereafter named AdrB3^SMC^). In contrast to AdrB3^EC^ mice, SMC restoration of β3-AR failed to have any significant effect on RVSP and Fulton’s index compared to AdrB3^KO^ mice (Sup. Fig. 3B-G). Our results also show that hypoxia increases SMC proliferation in both AdrB3^KO^ and AdrB3^SMC^ mice, showing a tendency towards less muscularization in AdrB3^SMC^ mice (Sup. Fig. 3H-K). Additionally, restoration of β3-AR expression in ECs tends to decrease ectopic expression levels of vWF in capillaries (Fig. 2L-M) and increase levels of nitrate in mice exposed to hypoxia (Fig. 2N) thus evidencing a direct protection over endothelium, leading to indirect effects over SMC.

**Figure 2.**
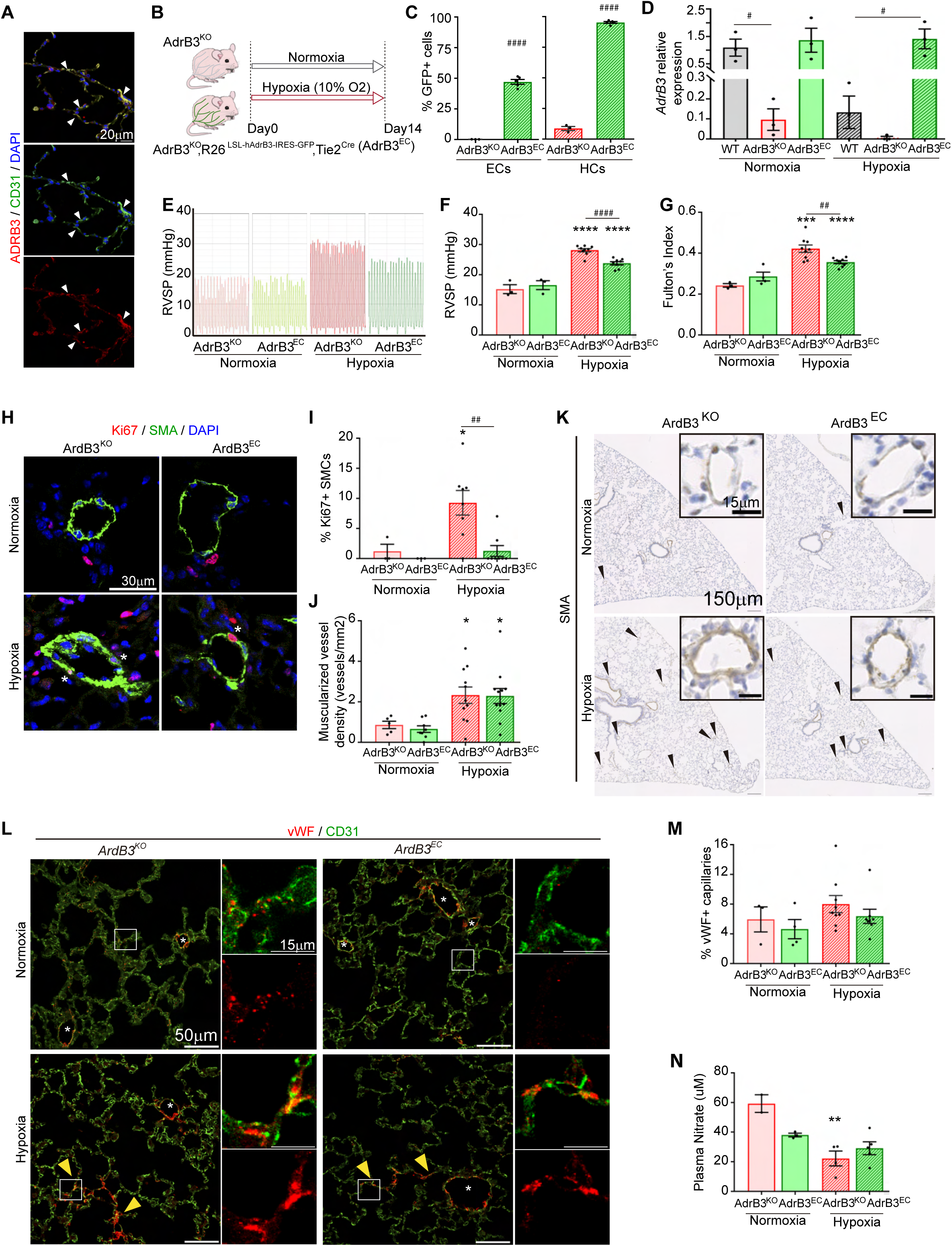
Restoration of endothelial β3-AR ameliorates the hypoxia-induced pulmonary hypertension phenotype in knock out mice. **(A)** Representative confocal images of β3-AR (red) and CD31 (green) immunostaining in mouse lung sections. **(B)** Hypoxia-induced PH experimental setup. **(C)** Quantification of frequency of transgene expression (R26^-LSL-hΒ3-AR-IRES-GFP^) in endothelial cells (ECs) and hematopoietic cells (HCs) in lungs of normoxic AdrB3^KO^ and AdrB3^EC^ mice. **(D)** β3-AR expression levels in lung ECs isolated by FACS from normoxic and hypoxic WT, AdrB3^KO^ and AdrB3^EC^ mice. **(E, F)** Representative recording profiles and quantification of RVSP. **(G)** Fulton’s index reflecting RV hypertrophy. **(H)** Representative confocal images of Ki67 (red) and SMA (green) immunostaining in mouse lung sections. Asterisks indicate proliferative Ki67^+^ SMCs. **(I)** Quantification of proliferating lung SMCs. **(J)** Quantification of density of fully muscularized arterioles in lung sections. **(K)** Representative images of mouse lung sections with immunostained for SMA (smooth muscle actin). Arrowheads indicate fully muscularized arterioles with less than 30µm diameter. Inserts depict representative arterioles. **(L)** Representative confocal images of vWF (red) and CD31 (green) immunostaining in mouse lung sections. Asterisks indicate arterioles, where vWF expression is observed in both normoxic and hypoxic conditions and yellow arrowheads indicate ectopic expression of vWF in capillaries. Insets of capillary regions are shown to better appreciate ectopic vWF expression. **(M)** Quantification of percentage of capillary bed with vWF expression. **(N)** Quantification of circulating nitrate as a readout of nitric oxide. All data are represented as mean ± s.e.m. Each dot in charts represents an individual patient/mouse. Two-tailed, unpaired t-test (B) and two-way ANOVA with Sidak’s multiple comparisons test for remaining charts. Asterisks indicate p-values between normoxic and hypoxic mice of the same genotype and hashtags indicate p-values between genotypes. *p<0.05, **p<0.01, ****p<0.001, #p <0.05, ##p<0.01, ####p<0.001.

**Figure 3.**
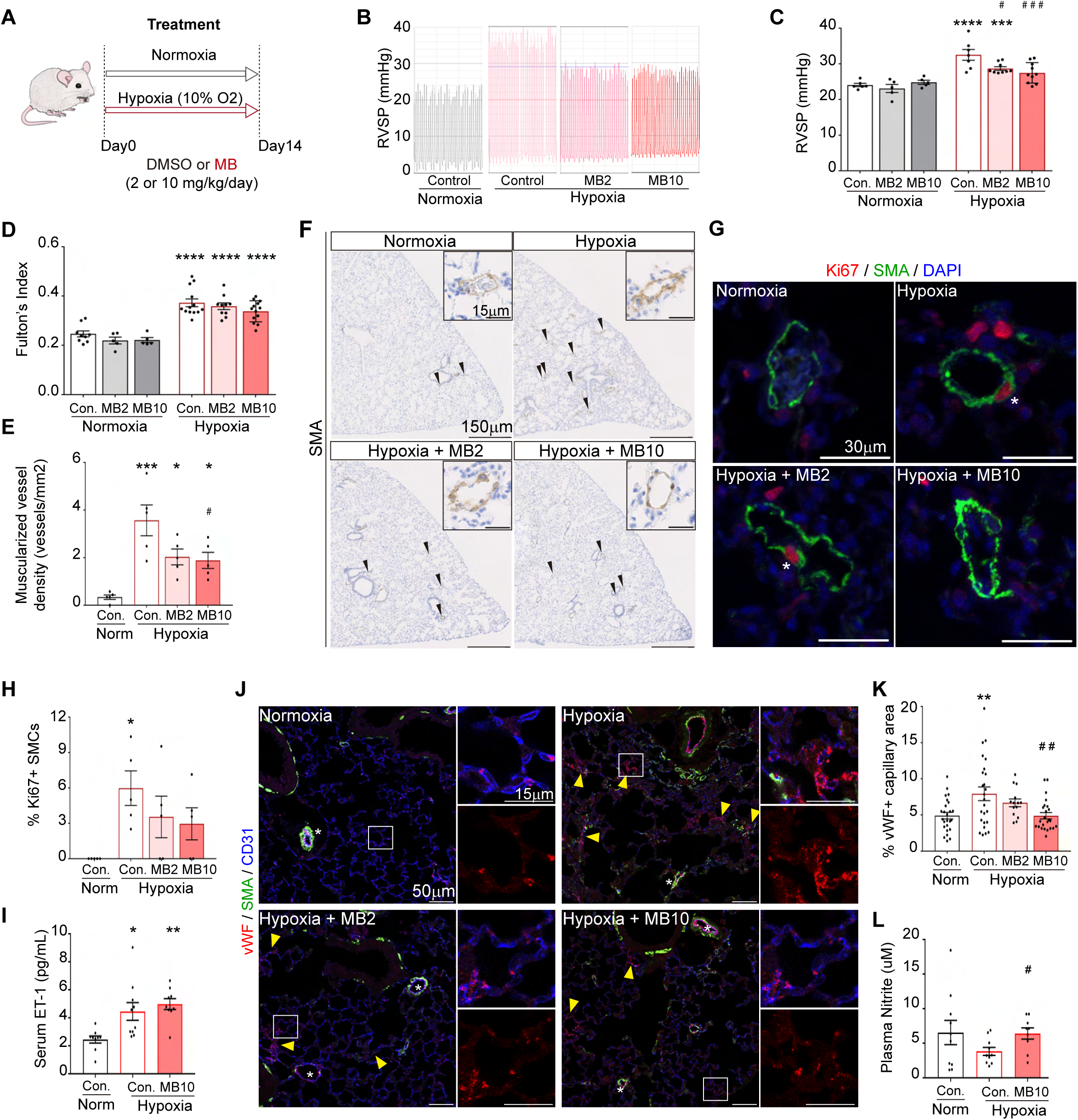
Mirabegron, a β3-AR agonist, prevents hypoxia-induced pulmonary hypertension in mice. **(A)** Hypoxia-induced pulmonary hypertension experimental setup and mirabegron or vehicle treatment scheme. **(B, C)** Representative recording profiles and quantification of RVSP. **(D)** Fulton’s index reflecting RV hypertrophy. **(E)** Quantification of density of fully muscularized arterioles in lung sections. **(F)** Representative images of mouse lung sections with immunostained for SMA (smooth muscle actin). Arrowheads indicate fully muscularized arterioles with less than 30µm diameter. Inserts depict representative arterioles. **(G)** Representative confocal images of Ki67 (red) and SMA (green) immunostaining in mouse lung sections. Asterisks indicate proliferative Ki67^+^ SMCs. **(H)** Quantification of proliferating lung SMCs. **(I)** Quantification of circulating ET-1 levels. **(J)** Representative confocal images of vWF (red), SMA (green) and CD31 (blue) immunostaining in mouse lung sections. Asterisks indicate arterioles, where vWF expression is observed in both normoxic and hypoxic conditions and yellow arrowheads indicate ectopic expression of vWF in capillaries. Insets of capillary regions are shown to better appreciate ectopic vWF expression. **(K)** Quantification of percentage of capillary bed with vWF expression. **(L)** Quantification of circulating nitrite as a readout of nitric oxide. All data are represented as mean ± s.e.m. Each dot in charts represents an individual patient/mouse. Two-way ANOVA with Sidak’s multiple comparisons test (C, D). One-way ANOVA with Dunnett’s multiple test comparisons for remaining charts. Asterisks indicate p-values between normoxic and hypoxic mice and hashtags indicate p-values between different treatments. *p<0.05, **p<0.01, ***p<0.005, ****p<0.001, #p <0.05, ##p<0.01, ###p<0.005.

### β3-AR activation prevents hypoxia-induced PH in mice in a dose-dependent manner

In order to confirm β3-AR as a potential therapeutic target in PH, we treated WT mice during the 2 weeks of exposure to chronic hypoxia with the β3-AR agonist mirabegron (MB) at two distinct doses (2mg/kg/day and 10mg/kg/day; hereafter named MB2 and MB10, respectively), or vehicle (DMSO 1%) in drinking water (Fig. 3A). Treatment with mirabegron attenuated most hypoxia-induced alterations in hemodynamic and histological parameters (Fig. 3B-J). A mirabegron dose-dependent decrease was observed in RVSP (32.51 ± 1.50 mmHg vs 28.71 ± 0.49 mmHg – p=0.013 – or vs 27.47 ± 0.91 mmHg – p=0.0007 – for DMSO, MB2 or MB10-treated mice, respectively) (Fig. 3B-C) and in RV hypertrophy (Fulton’s index: 0.373 ± 0.016 vs 0.359 ± 0.014, or vs 0.3379 ± 0.012 for DMSO, MB2 or MB10-treated mice) (Fig. 3D). Mirabegron did not have any effect in normoxic mice, which is in accordance with the absence of a role of β3-AR in normoxic conditions (see Fig. 1B). Histological analysis of the lung vasculature (Fig. 3E-H) confirmed the protective role of migrabegron-stimulated β3-AR signalling, as evidenced by a 50% reduction in the frequency of remodelled vessels (3.57 ± 0.65 vessels/mm^2^ vs 2.03 ± 0.33 vessels/mm^2^ – p=0.041 – or vs 1.883 ± 0.34 vessels/mm^2^ – p=0.025 – for DMSO, MB2 or MB10-treated mice, respectively) (Fig. 3E-F), and SMC proliferation compared to untreated mice (5.98 ± 1.46% vs 3.54 ± 1.76% vs 2.96 ± 1.35% for DMSO, MB2 or MB10-treated mice, respectively) (Fig. 3G-H). Endothelial dysfunction was assessed by confocal analysis of vWF immunostaining of lung sections and evaluation of circulating ET-1 and NO production. Although β3-AR-stimulation appeared to have no effect of circulating ET-1 levels (Fig. 3I), mirabegron-treated mice showed a significant decrease in vWF-expressing capillary ECs (7.93 ± 0.94% vs 6.675 ± 0.54% vs 4.886 ± 0.44% for DMSO, MB2 or MB10-treated mice, respectively; p=0.003 for MB10-treated mice) (Fig. 3J-K) and normalized circulating NO levels (Fig. 3L).

We also assessed cardiac function through echocardiography and obtained clear trends in several parameters that indicate that the RV size and function are better preserved in hypoxic mice treated with MB, than non-treated mice (Sup. Fig. 4A-G). Importantly, mirabegron attenuated the RV and PA dilatation induced by chronic hypoxia. In addition, hypoxia-associated alterations in pulmonary valve velocity-time integral (PV-VTI), pulmonary artery acceleration time (PAAT), tricuspid annular plane systolic excursion (TAPSE), and RV outflow tract (RVOT) were markedly reduced and, in some cases, restored towards normoxic values following mirabegron treatment. Additionally, and according to well-known cardiac metabolic alterations in PH, we observed that the increased GLUT1 protein expression in cardiomyocytes of mice with hypoxia-induced PH (0.45 ± 0.20% and 3.44 ± 0.81% in normoxic and hypoxic WT mice, respectively; p=0.013) (Sup. Fig. 4H-I) was attenuated following treatment with the higher dose of mirabegron.

**Figure 4.**
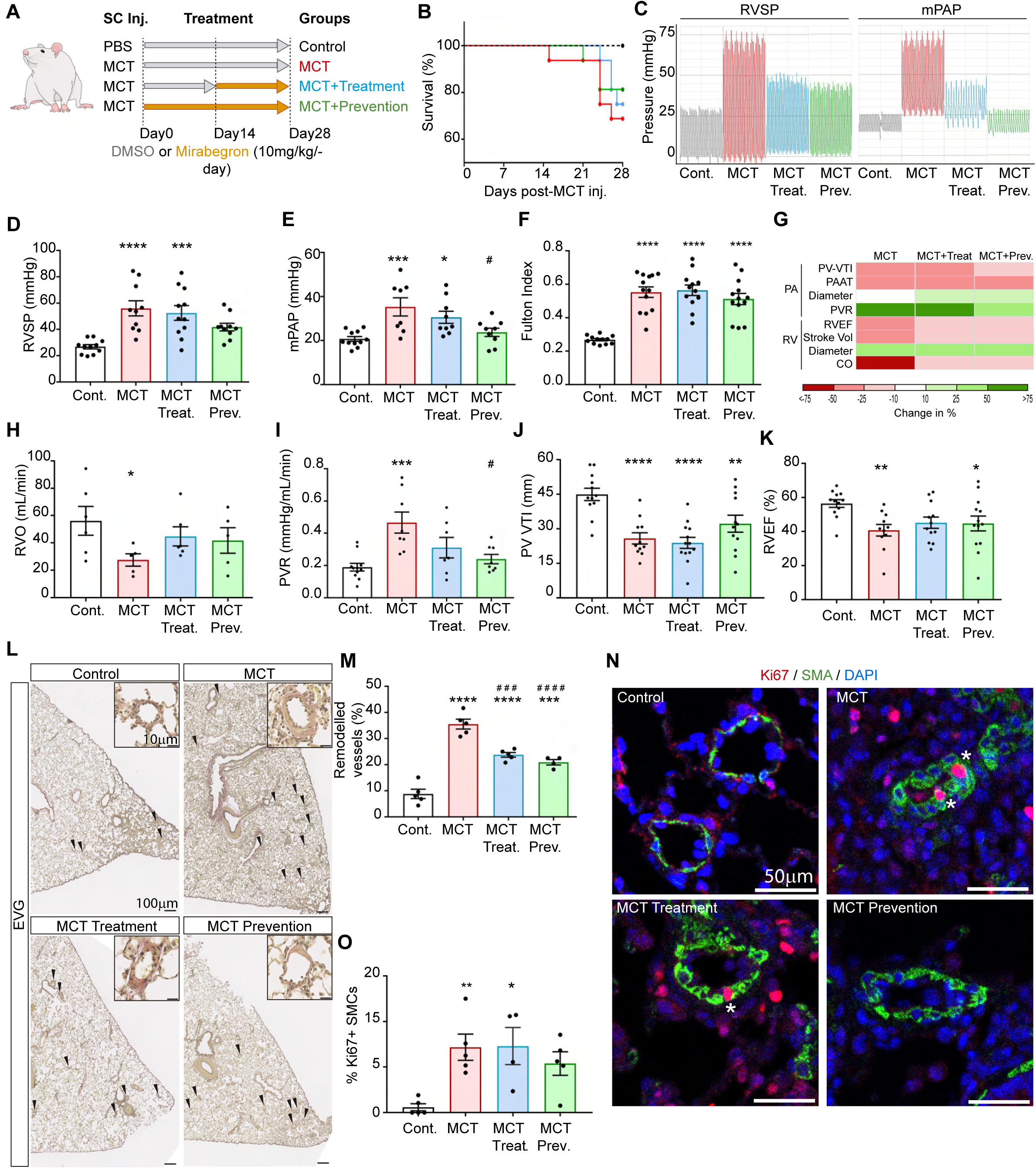
Mirabegron prevents MCT-induced pulmonary hypertension in rats. **(A)** MCT-induced pulmonary hypertension experimental setup and mirabegron/vehicle treatment scheme. **(B)** Survival curve of control or rats with MCT-induced pulmonary hypertension and subjected to different treatment schemes. **(C-E)** Representative recording profiles and quantification of RVSP and mPAP. **(F)** Fulton’s index reflecting RV hypertrophy. **(G)** Heat map representing change in percentage of several parameters determined by echocardiography of control or rats with MCT-induced pulmonary hypertension and subjected to different treatment schemes. **(H-K)** Charts with the quantification of parameters indicated in (G). **(L)** Representative images of rat lung sections with stained for EVG (Elastic Van Gieson). Arrowheads indicate remodelled arterioles with less than 100µm diameter. Inserts depict representative arterioles. **(M)** Quantification of density of remodelled arterioles in lung sections. **(N)** Representative confocal images of Ki67 (red) and SMA (green) immunostaining in rat lung sections. Asterisks indicate proliferative Ki67+ SMCs. **(O)** Quantification of proliferating lung SMCs. All data are represented as mean ± s.e.m. Each dot in charts represents an individual patient/mouse. One-way ANOVA with Dunnett’s multiple test comparisons. Asterisks indicate p-values between normoxic and hypoxic mice and hashtags indicate p-values between different treatments. *p<0.05, **p<0.01, ***p<0.005, ****p<0.001, #p <0.05, ###p<0.005, ####p<0.001.

### Mirabegron reduces pulmonary artery pressure and preserves cardiac function in the MCT-induced PH rat model

In order to rigorously validate the potential therapeutic value of the β3-AR-agonist mirabegron, we evaluated its effect in another pre-capillary PH model, the widely extended MCT-induced PAH in rats, where hypoxia is not the main driver of the disease. We randomized rats into 4 groups: control, diseased, treatment and prevention groups. Control rats received a subcutaneous injection of saline solution, while the remaining groups received a subcutaneous injection of MCT. The diseased group received no treatment, the treatment group received mirabegron (10mg/kg/day as the most effective doses in mice) at Day14, after the onset of the disease, and the prevention group was given mirabegron from Day0 after subcutaneous injection of MCT (Fig. 4A). The survival rate varied between the different groups with 32% of the rats of the MCT group having succumbed to the disease by Day28 (p=0.038), while the treatment and prevention groups presented only 25% (p=0.068) and 18% (p=0.120) of death rate, respectively (Fig. 4B). Mirabegron restored relevant hemodynamic and structural parameters, including RVSP, mPAP, PVR and pulmonary arteriole remodelling (Fig. 4C–O). Interestingly, RV functional parameters such as RV cardiac output (RVO) or RV Ejection Fraction (RVEF%) among others were recovered upon treatment with mirabegron (Fig. 4G-K, Supp. Table 2). The effects were observed in both treatment and prevention protocols, although they were generally more pronounced when mirabegron was administered from disease onset. Altogether, these results further support the potential therapeutic use of mirabegron in PH not only preventing but also being able to restore anatomical and functional alterations of the lung vasculature and the RV, even when the disease is already established.

### Endothelial β3-AR activity prevents SMC proliferation in a NO-dependent manner

Both loss- and gain-of-function genetic manipulation, as well as pharmacological stimulation of β3-AR in the *in vivo* experiments presented above suggest that β3-AR regulates EC dysfunction and SMC proliferation in mice exposed to hypoxia. In order to further explore the cellular and molecular mechanisms of β3-AR in the hypoxic lung vasculature, we checked *in vitro* reproducibility by using HPAECs and HPASMCs. We first assessed the non-cell autonomous role of endothelial β3-AR on HPASMC proliferation in hypoxic conditions, by culturing HPASMCs in hypoxia with conditioned medium from hypoxic HPAECs cultured in the presence or absence of mirabegron (Fig. 5A). We observed a 50% decrease in HPASMC proliferation when treated with conditioned medium of HPAECs treated with mirabegron. This effect of mirabegron was abrogated when HPAECs were also treated with L-NAME, a non-selective inhibitor of NOS enzymes, suggesting that endothelial β3-AR inhibits SMC proliferation via eNOS (Fig. 5B-C). In accordance with this, cAMP, a downstream target of β3-AR signalling and an activator of eNOS is increased in HPAECs (Fig. 5D-E), and supernatant of these same cells induced an increase in cGMP in HPASMCs (Fig. 5F). This strongly supports that activation of endothelial β3-AR inhibits SMC proliferation through the cAMP-eNOS-NO-cGMP axis. Concurrently, pharmacological activation of β3-AR ceases to have any beneficial effect on both pulmonary artery relaxation (Fig. 5I-J) and PH hemodynamic and morphological variables in NOS3^KO^ mice exposed to hypoxia (Fig. 5K-R).

**Figure 5.**
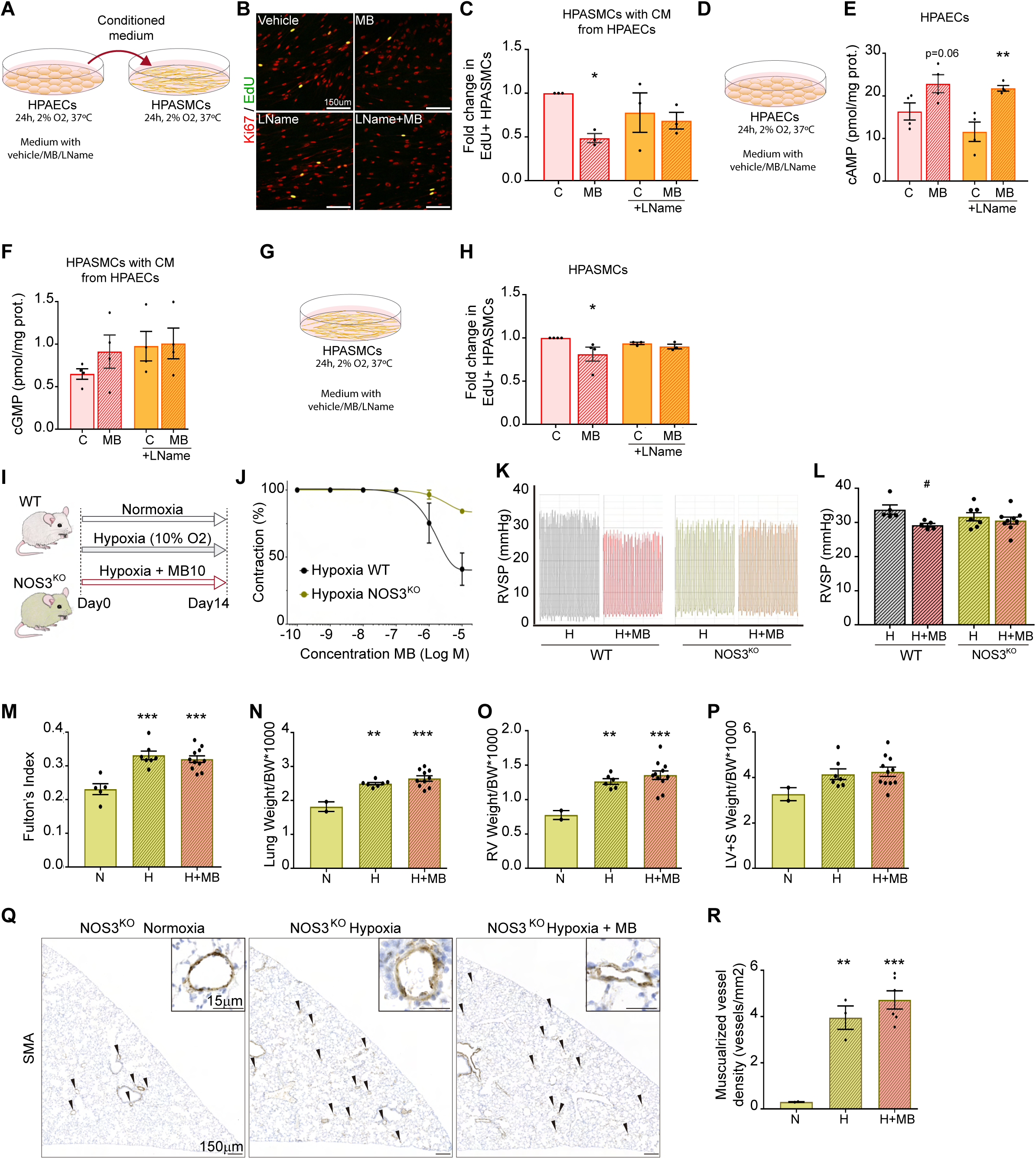
Endothelial β3-AR activity prevents SMC proliferation in a nitric oxide-dependent manner. **(A)** Experimental setup to study role of endothelial β3-AR on SMC proliferation. HPAECs were incubated 24 hours in a hypoxia chamber (2% O_2_) with medium supplemented with vehicle, mirabegron (10^-7^M) and/or L-Name (100µM). Conditioned medium of these cells was placed on HPASMCs, which were then incubated 24hrs in a hypoxia chamber (2% O_2_). HPASMC proliferation was assessed by adding EdU to medium 4 hours before fixing. **(B)** Representative confocal images of HPASMCs immunostained for Ki67 (red) and EdU (green). Double labelled cells are yellow. **(C)** Quantification of fold change in the percentage of proliferative HPASMCs. **(D)** Experimental setup to study role of Β3-AR on endothelial cells. HPAECs were incubated 24 hours in a hypoxia chamber (2% O_2_) with medium supplemented with vehicle, mirabegron (10^-7^M) and or L-Name (100µM). **(E)** Quantification of cAMP levels in HPAECs. **(F)** Quantification of cGMP levels in HPASMCs treated with conditioned medium from HPAECS, as in (A). **(G)** Experimental setup to study role of β3-AR on HPASMCs. HPASMCs were incubated 24 hours in a hypoxia chamber (2% O_2_) with medium supplemented with vehicle, mirabegron (10^-7^M) and/or L-Name (100µM). **(H)** Quantification of fold change in the percentage of proliferative HPASMCs, stimulated as in (G). **(I)** Hypoxia-induced pulmonary hypertension experimental setup. **(J)** Relaxation curves of phenylephrine pre-contracted hypoxic WT (black line) or NOS3^KO^ (green) pulmonary arteries in a wired myograph with different concentrations of the selective β3-agonist, mirabegron. **(K-L)** Representative recording profiles and quantification of RVSP. **(M)** Fulton’s index reflecting RV hypertrophy. **(N)** Ratio between left lung lobe and body weight of normoxic and hypoxic NOS3^KO^ mice with or without mirabegron (10mg/kg·day) treatment. **(O)** Ratio between RV and body weight of normoxic and hypoxic *NOS3^KO^*mice with or without mirabegron treatment. **(P)** Ratio between left ventricle + septum and body weight of normoxic and hypoxic *NOS3^KO^*mice with or without mirabegron treatment. **(Q)** Representative images of mouse lung sections immunostained for SMA. Arrowheads indicate remodelled arterioles with less than 30µm diameter. Inserts depict representative arterioles. **(R)** Quantification of density of remodelled arterioles in lung sections. All data are represented as mean ± s.e.m. Each dot represents an individual experiment or mouse. Two-way ANOVA with Sidak’s multiple test comparison (C, E, F, H, L) and One-way ANOVA with Tukey’s multiple test comparisons (M-P, R). Asterisks indicate p-values. *p<0.05, **p<0.01, ***p<0.005.

The cell autonomous role of β3-AR in HPASMC proliferation in hypoxic conditions was also assessed by directly culturing these in the presence or absence of mirabegron (Fig. 5G). Although our *in vivo* data indicated that reestablishment of β3-AR expression in SMCs does not ameliorate the PH phenotype of AdrB3^KO^ mice, including SMC proliferation (Sup. Fig. 3J-K), we observed a slight (18%) but significant decrease in HPASMC proliferation upon treatment with mirabegron (Fig. 5H). Although mirabegron exerted a modest but significant anti-proliferative effect in HPASMCs, the overall findings suggest that direct effects on SMCs contribute only partially to the protective actions of β3-AR activation, which appear to be predominantly mediated through the pulmonary endothelium.

### β3-AR stimulation inhibits hypoxia-induced ROS production and mitochondrial dysfunction in ECs

To assess endothelial dysfunction, we analysed several indicators of cellular stress *in vitro*: formation of stress fibres, cellular ROS production and mitochondrial morphology and function (Fig. 6). HPAECs subjected to hypoxia presented a significant increase in stress fibre formation which was attenuated by 25% with mirabegron (Fig. 6A, B). Cellular ROS production was increased in hypoxic HPAECs and β3-AR stimulation with mirabegron dramatically decreased this production (Fig. 6C, D). Hypoxia-induced mitochondrial fragmentation was also reduced with mirabegron (Fig. 6C, E). Although the effect of mirabegron on mitochondrial fragmentation was most pronounced at early timepoints (2 hours), analysis of transmission electron microscopy (TEM) images shows that cells treated with mirabegron continue to have better-preserved elongated mitochondria at later timepoints (Fig. 6F-I). When exposed to hypoxia, a clear shift in the distribution frequencies of mitochondria area is observed (Fig. 6G, H), with mitochondria from hypoxic cells presenting on average a 50% reduction in size when compared to mitochondria in normoxic cells. However, when cells are treated with mirabegron, this reduction in mitochondria size is significantly minimized (Fig. 6I), suggesting that mitochondrial fission is attenuated by the β3-AR pathway. To assess if mitochondrial functionality is also better preserved in hypoxic HPAECs treated with mirabegron, we evaluated mitochondrial respiration (Fig. 6K-O). Although overall mitochondrial content of HPAECs, assed by TOMM20 protein expression, does not change with hypoxia nor with mirabegron treatment (Fig. 6J), mitochondrial respiration is increased in mirabegron-treated cells (Fig. 6K-O) indicating a shift towards healthier ECs.

**Figure 6.**
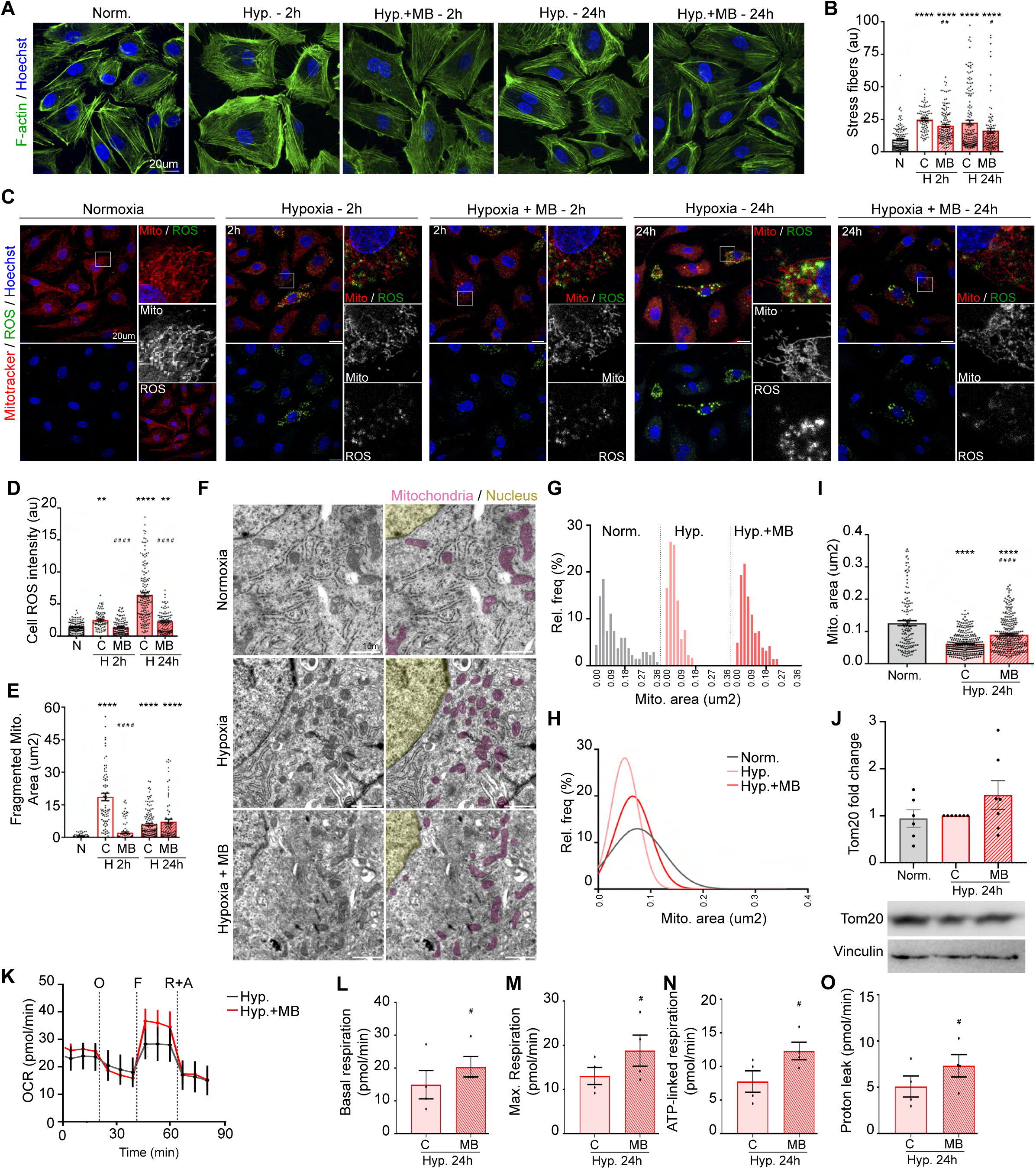
β3-AR stimulation inhibits hypoxia-induced ROS production and mitochondrial dysfunction in endothelial cells. **(A)** Representative confocal images of HPAECs exposed to hypoxia (2% O_2_) for 2 hour and 24 hours, in the presence or not of mirabegron (10^-7^M), and stained for F-actin (green) and Hoechst (blue). **(B)** Quantification of stress fibers from (A). **(C)** Representative confocal images of HPAECs exposed to hypoxia (2% O_2_) for 2 hour and 24 hours, in the presence or not of mirabegron (10^-7^M), labelled for mitochondria (MitoTracker, red) and cellular ROS (CellRox, green) and Hoechst (blue). **(D)** Quantification of mean intensity of cellular ROS. **(E)** Quantification of area occupied by fragmented mitochondria. **(F)** Representative TEM images of HPAECs exposed to normoxia or hypoxia (2% O_2_) for 24 hours, in the presence or not of mirabegron (10^-7^M). Images on the right are pseudocoloured to highlight nuclei (yellow) and mitochondria (pink). **(G-H)** Distribution histogram of mitochondrial sizes presents in each condition in (E). **(I)** Quantification of mitochondria area of HPAECs exposed to normoxia or hypoxia (with or without mirabegron). **(J)** Western blot and quantification of fold change of Tom20, a mitochondrial marker. **(K)** Oxygen consumption rate (OCR) of HPAECs after exposure to hypoxia (2% O_2_) during 24 hours, in the presence or not of mirabegron (10^-7^M). **(L-O)** Quantification of several parameters of mitochondrial respiration: basal respiration (L), maximal respiration (M), ATP-linked respiration (N) and proton leak (O). All data are represented as mean ± s.e.m. Each dot represents individual cells of 5-7 independent experiments (B, D; E) or individual experiments (I, J, L-O). One-way ANOVA with Dunnett’s multiple test comparisons (B, D, E), One-way ANOVA with Tukey’s multiple test comparisons (I-J), paired, two-tailed t-test (L-O). Asterisks indicate p-values in relation to normoxic conditions and hashtags indicate p-value in relation to cells not treated with mirabegron. **p<0.01, ****p<0.001, #p<0.05, ##p<0.01, ####p<0.001.

### β3-AR acts through an eNOS-UCP2 axis to balance ROS production and mitochondria respiration

Since the β3-AR activity has been previously linked to mitochondrial uncoupling proteins (UCP) and cell metabolism, we checked whether UCP1 or UCP2 levels were altered in our *in vivo* models. While UCP1 was undetectable in lung lysate, UCP2 was present both in normoxia and hypoxia. Importantly, in the absence of β3-AR, UCP2 was downregulated in hypoxic conditions (Fig. 7A), while mice treated with mirabegron presented increased UCP2 protein levels (Fig. 7B). Since UCPs are regulated by β3-AR at the transcriptional level via cAMP-activated PKA in other context,^25,31^ we questioned if UCP2 could also be regulated by β3-AR in the hypoxic pulmonary endothelium. For this purpose, we checked *ucp2* expression in isolated ECs by FACS from normoxic and hypoxic lungs of control, AdrB3^KO^ and mirabegron-treated mice. We did not observe any regulation of *ucp2* expression by either the loss or stimulation of β3-AR (Fig. 7C, D), suggesting that β3-AR-driven changes in UCP2 protein levels occur post-transcriptionally. As we showed above that *in vivo* stimulation of β3-AR ceases to have any beneficial effect on hypoxic NOS3^KO^ mice, we checked if β3-AR-driven regulation of UCP2 could be eNOS dependent. Indeed, in the absence of eNOS and under hypoxic conditions, stimulation of β3-AR no longer leads to an upregulation of UCP2 (Fig. 7E).

**Figure 7.**
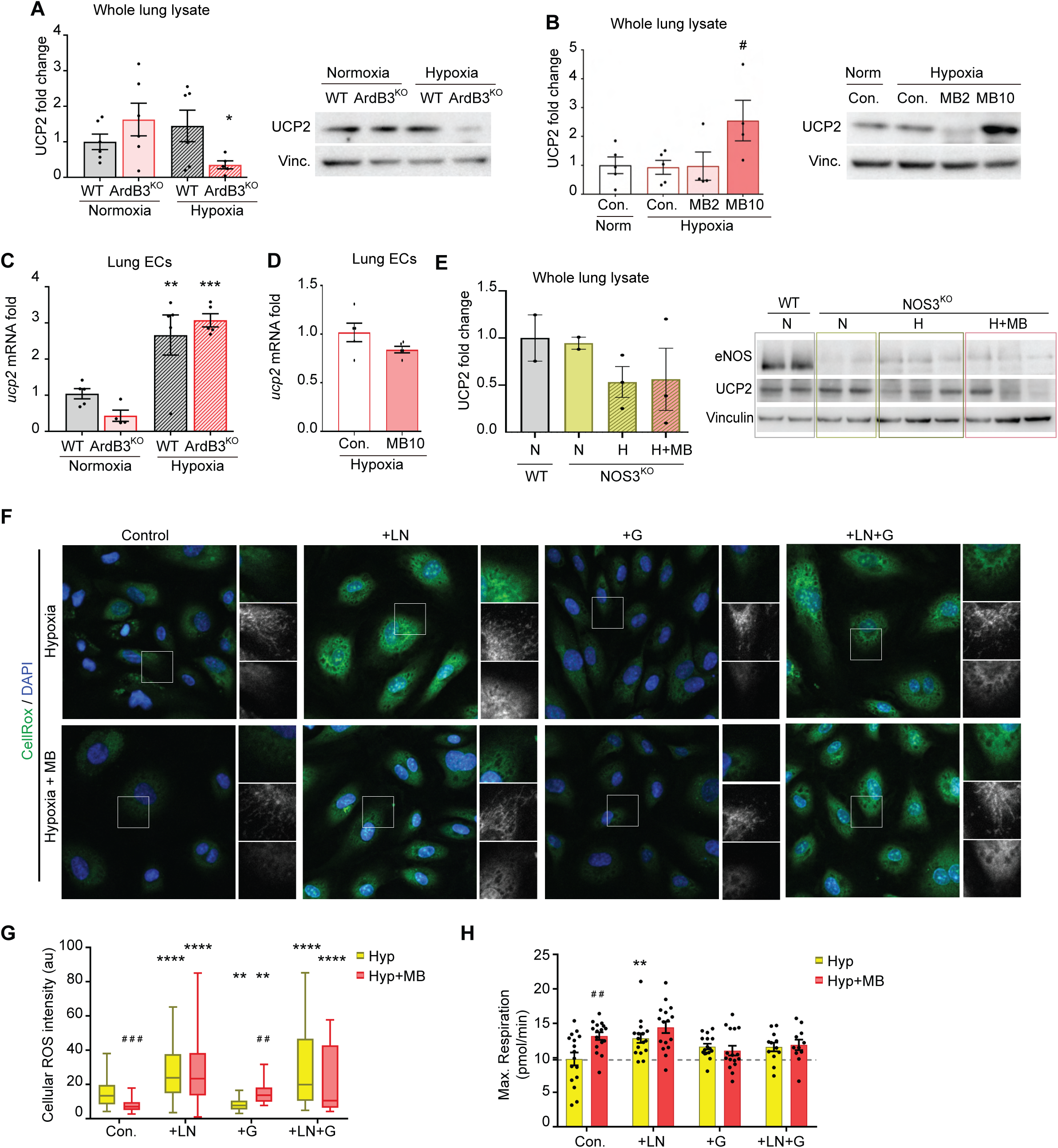
β3-AR acts through an eNOS-UCP2 axis to balance ROS production and mitochondria respiration. **(A)** Quantification and representative western blot of UCP2 in total lung lysate of WT or AdrB3^KO^ mice exposed to normoxia or hypoxia (as in Fig. 1A). **(B)** Quantification and representative western blot of UCP2 in total lung lysate of normoxia or hypoxia-exposed mice treated with mirabegron or vehicle (as in Fig. 3A). **(C)** mRNA expression levels of ucp2 in endothelial cells isolated by FACS from lungs of WT or AdrB3^KO^ mice exposed to normoxia or hypoxia (as in Fig. 1A). **(D)** mRNA expression levels of ucp2 in endothelial cells isolated by FACS from lungs of hypoxia-exposed mice treated with vehicle or mirabegron at 10mg/kg/day (as in Fig. 3A). **(E)** Quantification and representative western blot of UCP2 in total lung lysate of WT or NOS3^KO^ mice exposed to normoxia or hypoxia, with or without mirabegron treatment at 10mg/kg/day (as in Fig. 5I). **(F)** Representative confocal images of HPAECS incubated for 24 hours in hypoxia (2% O_2_) in the presence or absence of mirabegron (10^-7^M, MB), L-NAME (100µM; LN) or Genipin (10µM, G). ROS was detected by incubation cells with CellROX Green for 30 minutes, 37°C prior to fixation and imaging. **(G)** Quantification of mean intensity of cellular ROS in (F). **(H)** Quantification maximal respiration of HPAECS incubated for 24 hours in hypoxia (2% O_2_) in the presence or absence of mirabegron (10^-7^M, MB), L-NAME (100µM; LN) or genipin (10µM, G). All data are represented as mean ± s.e.m. Each dot represents mice (A-E) or 6-8 replicates for 2 independent experiments (H). Two-way ANOVA with Sidak’s multiple test comparison (A, C, G, H), One-way ANOVA with Tukey’s multiple test comparisons (B), unpaired, two-tailed t-test (D). Asterisks indicate p-values in relation to normoxic (A, C; E) or hypoxic with no inhibitors (G, H) conditions and hashtags indicate p-value in relation to cells not treated with mirabegron. *p<0.05, **p<0.01, ***p<0.005, ****p<0.001, #p<0.05, ##p<0.01, ###p<0.005.

Finally, to better understand whether the protective role of β3-AR in ECs from ROS is mediated via eNOS and/or UCP2, we assessed the effect of β3-AR stimulation on cellular ROS production and mitochondrial respiration when inhibiting eNOS and/or UCP2 (Fig. 7F, G). As we already shown, β3-AR activation leads to reduced ROS production and increased mitochondrial respiration (also shown in Fig. 7G, H), which we attribute to the better preservation of mitochondria due to NO-mediated ROS scavenging. In this line, L-NAME-mediated inhibition of eNOS leads to both a dramatic increase in ROS production and enhanced mitochondrial respiration (Fig. 7G, H). These effects are in accordance with the anti-oxidant role of NO^32^ and to its inhibitory role on cytochrome c oxidase and consequently mitochondrial respiration.^33^ As it is shown in Fig. 7F-H, stimulation of β3-AR in the presence of L-NAME is, as expected, unable to revert this increase in ROS production and has no additional effect on mitochondrial respiration, as β3-AR acts through eNOS. Additionally, UCP2, which seems to be downstream of the β3-AR-eNOS axis was inhibited with genipin. Whereas UCP2 inhibition per se did not yield major effects on ROS production or mitochondrial respiration, it either inhibited or inversed β3-AR effects. On these conditions, stimulation of β3-AR with mirabegron no longer led to increased mitochondrial respiration and ROS production was even slightly increased, albeit not to the levels observed upon L-NAME treatment, as UCP2 inhibition does not affect NO production. Lastly, combined inhibition of both eNOS and UCP2, lead to the same outcome as eNOS inhibition alone, further supporting the hypothesis that UCP2 acts downstream of the β3-AR-eNOS axis.

## DISCUSSION

In this study we have confirmed a pivotal role of β3-AR in regulating vascular endothelial function and fate thus leading to preventing development of pre-capillary forms of PH. We have demonstrated for the first time the presence of β3-AR in human pulmonary endothelium of patients with severe COPD, where low oxygen is present. This finding correlates with our results in hypoxia-induce PH mice where β3-AR appears to be overexpressed, probably as a compensatory mechanism against the increase of vascular tone characteristic of PH. In this line, we have demonstrated how the deletion of the β3-AR gene worsens the PH phenotype, while specific restoration of the receptor in the EC recovers hemodynamic function and indirectly decreases SMC proliferation in our mouse model. Our findings also displayed that the administration of the β3-agonist mirabegron decreased vascular remodeling, conserved hemodynamic variables, and preserved the RV function both in hypoxia- and monocrotaline-induced PH mice and rats respectively. Furthermore, our studies in human cells and NOS3^KO^ mice demonstrate that mirabegron acts directly on ECs through eNOS/NO and UCP2 dependent mechanisms, reducing oxidative stress and preserving mitochondrial homeostasis. Overall, our findings identify β3-AR as a promising therapeutic target and mirabegron as a potential therapeutic strategy to prevent endothelial damage, vascular remodeling, and cardiovascular dysfunction in pre-capillary forms of PH.

Although β3-AR was originally described in the adipose tissue, it has also been described in cardiac and vascular cells, especially in pathologies such as systemic hypertension, myocardial infarction and HF, where it plays an important role in regulating vasodilation and mitigating cardiac injury.^20,23–25^ This particular adrenergic receptor, which is not present in healthy vascular and cardiac tissues, suffers an overexpression during pathologies.^20,34,35^ This phenomenon has been previously demonstrated by us and others and described as a compensatory mechanism due to lack of active β1-AR and β2-AR, that are common cardiovascular adrenergic receptors.^34,36,37^ However, this potential compensatory mechanism has also been demonstrated not to be enough to stop pathology because β3-AR is also the less sensitive to endogenous catecholamines among all adrenergic receptors.^20,35,37^ In this situation, either higher levels of expression or an activation by exogenous β3-agonist has demonstrated to be more effective. Additionally, in previous works where β3-AR was overexpressed using adeno-associated viral particles, it was observed a reduction in cardiac fibrosis and an improvement LV systolic function in mice. Overexpression of β3-AR in cardiomyocytes also showed a basal remodeling of the entire mitochondrial network, increasing mitochondrial biogenesis and reducing mitophagy.^25,38^ All these evidences, together with our results suggest that β3-AR could be a relevant target in PH.

In previous works done by others, β3-AR has been shown to be regulated by hypoxia in the context of retinal diseases^39,40^. Accordingly, we demonstrate that β3-AR in up-regulated in pulmonary endothelium from animal models of hypoxia-induced PH and COPD patients. This evidence led us to think that β3-AR could have a potential role in hypoxic related forms of PH. One of these forms, PH associated to lung diseases (group 3), is also originated in severe COPD.^41^ Patients with COPD present alveolar hypoxia and can develop PH depending on the severity of the disease and its stage.^42^ Several studies in patients with stage IV COPD showed a mPAP > 20 mmHg and the appearance of vascular lesions.^13,43^ Our results showed for the first time that β3-AR is overexpressed in the pulmonary endothelium of severe COPD patients, directly correlating with BODE index. Additionally, when we explored the activity of this receptor in mice subjected to hypoxia-induced PH, we found that isolated pulmonary arteries from mice subjected to chronic hypoxia reacted in a concentration-dependent manner upon stimulation with β3-agonist in *ex vivo* wire myograph experiments. However, control mice arteries did not relax, thus confirming that hypoxic conditions favor an up-regulation of β3-AR in pulmonary arteries. In parallel, AdrB3^KO^ mice subjected to chronic hypoxia developed an aggravated PH phenotype compared to WT controls, thus supporting the idea that β3-AR overexpression appears as an attempt of the organism to compensate the increase in vascular tone, as we and other suggested in other contexts before.

β3-AR could be expressed in different cell types such as ECs and SMCs, where it can be contributing to the health/disease balance in a different manner,^34^ however its presence in pulmonary arterioles remained unknown. Also, considering its potential protective effect we wanted to elucidate whether this activity came from β3-AR in ECs and/or SMCs. For this purpose, we crossbred our AdrB3^KO^ mice with conditional transgenic mice overexpressing human β3-AR depending on the particular Cre line presented. Therefore, we generated mice lacking β3-AR in the whole body and restored its expression either in EC or SMC. Our results demonstrated that only those mice in which we restored β3-AR levels in ECs had an attenuation of the PH phenotype, showing less RVSP, RV hypertrophy and vascular remodeling, and including recovery of damaged endothelium. These findings suggest that the protective effects of β3-AR in PH are driven largely by the pulmonary endothelium, underscoring the relevance of endothelial β3-AR signaling in the regulation of vascular remodeling and disease progression.

Since the use of β3-agonist has been previously proposed to be effective in animal models of PH,^21,44^ we decided to run preclinical trials with two of the most common animal models of the disease: 1) the chronic hypoxia induced PH model, where hypoxia is the main driver; and 2) the MCT induced PAH rat model, a widely studied model of disease that allow us to further explore other features of the disease in different time-frames. For this purpose, we selected mirabegron, which was approved by drug agencies for use in the treatment of overactive bladder.^45^ This drug emerges as a potential drug repurposing strategy which, in addition to its demonstrated effectivity, would facilitate its translation into humans. In fact, this drug has also been proposed for its use in advanced stages of HF^46^ and PH associated with LV failure.^47^ Our results demonstrated that even lower doses of mirabegron where effective to reduce RVSP, vascular remodeling and RV hypertrophy in pre-capillary forms of PH in mice and MCT-rats, in a dose-responded manner. Mirabegron was enough to recover animals with PH and also resulted in a healthier endothelium – evidenced by lowering tissular and circulating markers of endothelial damage – and regulated cardiac metabolism and function. Remarkably, even rats receiving the drug two weeks after the development of the disease were found in better condition than PH disease controls. Although the effect on RV hypertrophy was less evident, echocardiographic evaluation demonstrated improvements in CO, PVR, and RVEF, all of which are important prognostic markers in PH. Furthermore, consistent with our findings, mirabegron administration improved RV function in MCT-rats and group 2 PH patients^44,47^. These results demonstrate the ability of mirabegron to preserve pulmonary hemodynamics and cardiac function while limiting vascular remodeling. Collectively, these findings further support β3-AR activation as a promising therapeutic strategy for pre-capillary PH.

It is well known that endothelial β3-AR can induce relaxation through an eNOS-dependent mechanisms.^48^ However, the exact mechanism of β3-AR in PH pulmonary arteries was not yet elucidated. To further explore the cellular and molecular mechanism behind the positive effect of β3-AR activation by mirabegron, we use a set of experiments including both NOS3^KO^ mice *in vivo* and pharmacological tools *in vitro*. When agonist mirabegron was administered to mice under chronic hypoxia with genetic ablation for eNOS, no effect was seen in the PH phenotype. Also, NOS3^KO^ pulmonary arteries, as well as those from hypoxic AdrB3^KO^ mice were unable to vasodilate in response to mirabegron stimulation. These observations are consistent with the functional impairment of the endothelial eNOS/NO axis identified in AdrB3^KO^ mice, which display reduced NO production despite maintaining normal eNOS protein levels, and our *in vitro* results using L-NAME, which confirms the role of endothelial β3-AR/eNOS pathway as expected. However, in SMCs β3-AR works through another mechanism. In these cells, the receptor is coupled to Gs protein that leads to the activation of adenylate cyclase and the production of cAMP, which generates vasorelaxation through the cAMP/PKA pathway.^49,50^ Furthermore, in SMCs, cGMP plays a very important role in cell proliferation.^50^ As we also found a marked reduction in vascular remodeling and Ki67 markers in rodents treated with mirabegron, we wanted to clarify whether β3-AR had an effect directly to SMC or not. Our mice with restored β3-AR in SMC could not scope the effect of hypoxia and developed PH. In concordance, our *in vitro* experiments using HPASMC demonstrated that mirabegron did not have a major effect on cell proliferation when incubating directly with this cell type. However, culturing HPASMC with supplemented media from previously incubated with mirabegron HPAEC, resulted in a marked inhibition of proliferation. On the contrary, this effect was abrogated when using preconditioned media from ECs treated both with mirabegron and the eNOS inhibitor L-NAME, thus confirming that the effect exerted by the β3-agonist over SMC, was more of an indirect effect due to the NO production. In fact, under healthy conditions, ECs produce NO that diffuses towards SMCs activating sGC that converts GTP into cGMP favors the inhibition of cell proliferation.^7,38^ The fact that mirabegron had no effect on SMCs when culturing with a medium supplemented with ECs treated with an eNOS inhibitor, confirmed this mechanism. In addition, our results showed that the absence of β3-AR in mice aggravates vascular remodeling, however, its overexpression in ECs improves hemodynamic measures and decreases the proliferation of SMCs. Therefore, our results suggest that β3-AR acts mainly on the endothelium, inhibiting the proliferation of SMCs in an endothelial NO dependent manner, also contributing to reduce RV remodeling and improving hemodynamic measures.

To further explore the molecular mechanisms behind the protection of pulmonary endothelium, we performed mechanistic assays using HPAEC subjected to hypoxia. ECs under pathological conditions have been shown to have elevated stress actin fibers and higher ROS production.^51^ The activation of β3-AR with selective concentrations of mirabegron abrogated changes induced by hypoxia. β3-AR has been previously described to be connected to energy metabolism and mitochondrial fitness.^24,25,52^ Mitochondria are essential organelles for energy production, calcium homeostasis, redox signaling, and other pathological processes. Mitochondrial homeostasis depends on a balance in mitochondrial fusion and fission processes known as mitochondrial dynamics. Some pathological processes that generate hypoxia can alter this dynamic, increasing ROS and decreasing ATP production.^53^ In fact, HPAEC under hypoxic conditions presented a hyper-fragmentation of the mitochondrial network also coincident with an impaired respiration and higher ROS production. Activation of β3-AR prevented both the mitochondrial fragmentation and rise in ROS. Among all the proteins linked to mitochondrial activity, UCPs are mitochondrial transporters located in the mitochondrial membrane expressed ubiquitously in most cell types. Certain UCPs has been linked to β3-AR activity regarding cell metabolism and energy consumption.^25,54^ Among them, UCP2 stands by its ability to exert a protective effect against oxidative damage.^55^ The role of UCP2 has been recently discussed.^56^ While its most extended role has been assumed to be acting by reducing the electrical potential across the inner mitochondrial membrane, thereby reducing the production of superoxide radical in complex I of the mitochondrial electron transport chain,^56,57^ this mechanism remains controversial. On the other hand, we have recently demonstrated that UCP2 could also act as a metabolite carrier to regulate cell proliferation and cytosolic ROS.^56^ UCP2 has been also previously described to be involved in PH.^58^ In the absence of oxygen, UCP2 is downregulated thus as a normal response to lack of oxygen coincident with reduced mitochondrial respiration.^58^ Our results have shown that the activation of β3-AR through mirabegron induces an increase in the expression of UCP2, which in consequence could be partially responsible of the decreased ROS and subsequently in the EC survival. Additionally, we found an improvement in mitochondrial fitness, reduced fragmentation and increased respiration overall contributing to the normal functioning of the cell. In fact, mitochondria in ECs subjected to hypoxia are fragmented, unlike those from ECs treated with mirabegron, where mitochondrial fragmentation decreases. On the contrary, in AdrB3^KO^ mice we found a decrease in the expression of UCP2 suggesting that this protein would be regulated by β3-AR downstream pathway. Furthermore, in cells with pharmacological inhibition of UCP2 we observed that the administration of mirabegron did not decrease ROS levels and did not increase mitochondrial respiration, demonstrating the important role that UCP2 plays in the control of cell ROS and mitochondrial function.^56,59^ In addition to our findings, it has been previously described the link between β3-AR and the activation of AMPK.^60^ This activation of AMPK will likely follow the activation of the peroxisome proliferator-activated receptor gamma co-activator 1 alpha (PGC-1α), which in the nucleus increases the expression of the UCP2 protein, that generates a reduction in ROS and increases the bioavailability of NO,^25,61^ thus confirming our results.

Overall, our findings give strong evidence supporting a protective role of the endothelial β3-AR signalling pathway in pre-capillary forms of PH. Additionally, we demonstrated that β3-AR activation by mirabegron preserves endothelial function and modulates vascular remodeling through two distinct mechanisms: 1) by the activation of eNOS and NO production which in turns ameliorates SMC relaxation capabilities and regulates cell proliferation; 2) via upregulation of the UCP2 protein which can reduce ROS and protect mitochondrial structure and fitness. These changes would help to preserve the integrity of the endothelium, being able to correctly regulate vascular remodeling and vasodilation, to reduce both mPAP and PVR, and to improve cardiac function, which delays the progression of PH and the appearance of RV failure (Fig.8), suggesting mirabegron as a relevant emerging tool to protect pulmonary endothelium in those forms of PH where β3-AR will be present. In this regards, the use of mirabegron in PH associated with LV failure (group 2) has been recently investigated in a clinical trial (SPHERE-HF).^47^ Although this trial found positive results in the RVEF of patients treated with mirabegron, the primary endpoints (PVR) was unchanged. Therefore, mirabegron helps improve heart function as we have also seen in our mice and rat’s results; however, it may not be sufficiently helpful in CpcPH patients because of their advanced endothelial dysfunction and more severe vascular injury.^4,62^ PH associated with LV failure begins in advances steps of chronic HF as a consequence of a backward surge of pressure into the pulmonary circulation,^63^ which in turns induces an elevated pressure injury of the capillary wall (stress failure),^64^ thus weakening the integrity of the alveolar-capillary unit and altering endothelial integrity. As we have demonstrated herein, β3-AR activation in PH would mainly act via pulmonary endothelium. Therefore, the status of the intima layer could be essential to ensure the usefulness of this therapeutic strategy. Instead, patients with pre-capillary forms of PH – group 1 (PAH) and group 3 (CLD-PH) associated to lung disease and/or hypoxia – are patients with a different origin of the disease, so the pathology differs cellular and molecularly.^12,13^ These group of patients are of special interest since few specific therapies has been approved. More importantly, while currently approved therapies stand by the assumption of an existing endothelial injury acting over SMC layer, this is to the best of our knowledge, the first therapy to propose repairing and protecting the pulmonary endothelium. This not only suggests the relevance of EC layer in controlling pulmonary vascular homeostasis, but also suggests that therapies regulating cell healthy activity could be an interesting therapeutic option alone or in combination with other existing therapies in order to conquer a full vascular protection in PH patients.

**Figure 8.**
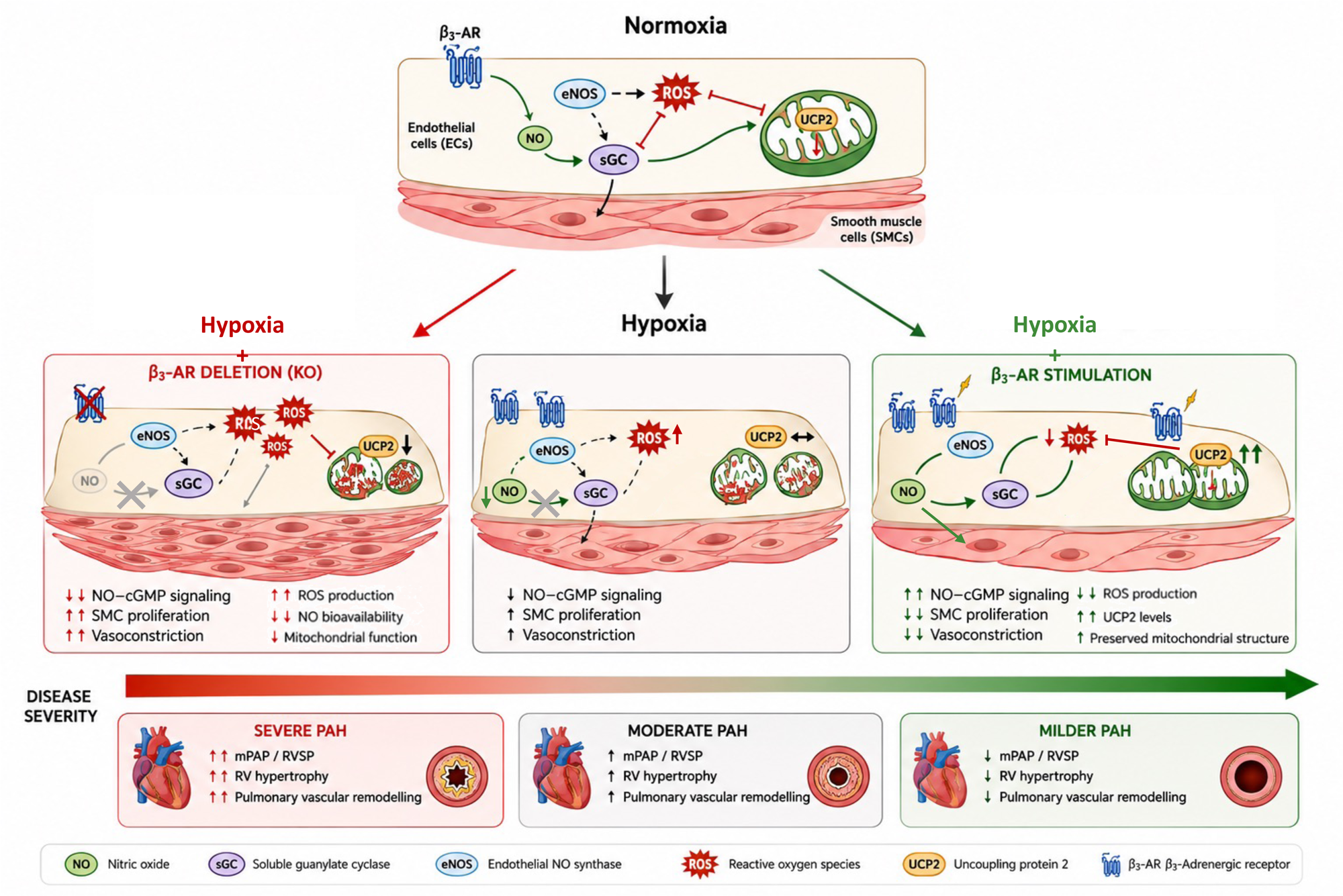
β3-AR as a key regulator of endothelial function and PH progression. Schematic summary of the effects of β3-AR signalling in pulmonary ECs and SMCs under normoxia, hypoxia with β3-AR deletion, and hypoxia with β3-AR stimulation. In normoxia, β3-AR expression is low and vascular homeostasis is maintained. Under hypoxia with β3-AR deletion, endothelial dysfunction is aggravated, with decreased eNOS activity, reduced NO bioavailability, increased ROS production, impaired mitochondrial function, and enhanced SMC proliferation and vasoconstriction, leading to severe pulmonary vascular remodeling, RV hypertrophy, and elevated RVSP. A similar pattern is observed in WT mice exposed to hypoxia, although the magnitude of these alterations is less pronounced than in β3-AR KO mice. In contrast, pharmacological activation of β3-AR with mirabegron preserves endothelial function by sustaining eNOS/NO signalling, upregulating UCP2, reducing ROS, and maintaining mitochondrial homeostasis. These protective effects limit SMC proliferation, promote vasodilation, and ameliorate pulmonary vascular remodeling and RV adaptation, resulting in an overall milder PH phenotype. Collectively, these findings support the therapeutic relevance of targeting β3-AR signalling and highlight the translational potential of mirabegron in pre-capillary forms of PH.

In conclusion, we herein demonstrate that endothelial β3-AR plays a relevant role controlling pulmonary vascular homeostasis and remodeling in PH, and its activation with mirabegron preserves the integrity and functionality of the endothelium, being able to correctly regulate vasodilation and cell proliferation, to improve hemodynamic variables and to maintain RV function in animal models of PH. In this incoming era when new therapies are approaching PH patients to regulate vascular remodeling, adjuvant strategies protecting pulmonary endothelium became necessary to act against the disease from a multiple perspective and help prevent adverse outcomes. These findings provide a compelling translational basis for modulating the β3-adrenergic pathway in PH, being the agonist mirabegron a potential drug repurposing strategy for the treatment of pre-capillary forms of PH.

## ACKNOWLEDGMENTS

Conditional β3-AR over-expressed (R26^LSL-Β3-AR-IRES-GFP^) mice were generated by Jose Luis de la Pompa’s laboratory at CNIC. The design of the graphical abstract was improved using ChatGPT 5.5. We thank the CNIC and CIB-CSIC Animal, Microscopy and Advance Imaging units for support.

