## Supplemental Material for "Endothelial β3-Adrenergic Receptor activation prevents pulmonary hypertension"

#### SUPPLEMENTAL METHODS

##### *Study design*

All experiments with animals conformed to EU Directive 2010/63EU and Recommendation 2007/526/EC, enforced in Spanish law under Real Decreto 53/2013. Animal protocols were approved by the local ethics committees and the Animal Protection Area of the Comunidad Autónoma de Madrid, the Universidad Autónoma de Madrid-CNIC (PROEX 112.5/20) and CIB-CSIC (PROEX 195.2/22), and conformed to ARRIVE guidelines 2.0. All animal experiments were conducted by authorized experienced researchers blinded to experimental groups.

##### *Animals model generation*

In order to evaluate the role of β3-AR in PH, β3-AR global knockout mice ($AdrB3^{KO}$)[1] were generated. To reestablish β3-AR expression specifically in ECs or SMCs, $AdrB3^{KO}$ mice were crossbreed with $AdrB3^{EC\text{-}OE}$ or $AdrB3^{SMC\text{-}OE}$ mice, respectively. $AdrB3^{EC\text{-}OE}$ or $AdrB3^{SMC\text{-}OE}$ mice were obtained by crossbreeding a mouse line that carries a β3-AR conditional over-expression allele ($R26^{LSL\text{-}B3\text{-}AR\text{-}IRES\text{-}GFP}$)[2,3] with an EC (Tie2-cre)[4] or SMC (Sm22-Cre)[5] Cre driver. $NOS3^{KO}$ mice were obtained from The Jackson Laboratory (B6.129P2-Nos3tm1Unc/J, stock No. 002684). Adult male C57BL6 WT, transgenic mice lines and Sprague-Dawley rats were used in this study. Animals were randomly allocated to the different experimental groups.

##### *Pulmonary Hypertension animal models*

*Hypoxia-induced pulmonary hypertension in mice:* male mice at 12 weeks of age (25-30g) were exposed to chronic hypoxia at 10% $O_2$ atmosphere inside a ventilated chamber, for 2 weeks. The hypoxic chamber was opened once a week to replenish food and water and to change the bedding. Control mice were kept in normoxic conditions, in the same animal room outside the hypoxic chamber. In randomized experiments, mice were administrated mirabegron (Santa Cruz Biotechnology, sc-211912) in drinking water at 2 or 10 mg/kg/day or with equal volume of vehicle only (DMSO). Drinking bottles were refreshed weekly. Echocardiographic analysis was performed in mice after 2 weeks of exposure to chronic hypoxia (Day 14). At endpoint, mice were anesthetized, animals underwent RVSP measurement through cardiac puncture, quickly euthanized and tissues collected.

*Rat monocrotaline-induced PH:* PH was induced by subcutaneous injection of monocrotaline (MCT; 60mg/kg; Sigma, C2401) in the inner hind leg of Sprague-Dawley male rats (240–300 g). Control rats were injected with saline solution and MCT-injected rats were randomized into 3 groups: MCT, MCT+Prevention and MCT+Treatment. The MCT+Prevention group received mirabegron (10 mg/kg/day) in drinking water from the day of MCT injection until week 4. In contrast, the MCT+Treatment group received mirabegron between weeks 3 and 4 after MCT injection. Moreover, control and MCT groups received drinking water with vehicle only (DMSO) and drinking bottles were refreshed every 3 days. Echocardiographic analysis was performed in rats at day 28 (4 weeks) after injection of MCT or saline solution. At endpoint, rats were anesthetized, hemodynamic measurements were taken, quickly euthanized and tissues collected.

***Hemodynamic measurements and tissue collection***

*Mouse RVSP measurement:* mice were deeply anesthetized intraperitoneally with a mix of medetomidine (1mg/kg) and ketamine (75mg/kg). Mice were placed in supine position over a heating pad to maintain body temperature at 37ºC, approximately, and skin from thorax was removed. RVSP was measured by closed-chest insertion of a 27 G catheter (Venofix A), coupled to a pressure transducer (Transpac IV), directly into the RV. Hemodynamic data was recorded using a Biopac MP36R System and Biopac's Aqcknowledge 4.1.0 software. For each mouse, at least 30 second of continuous and stable heartbeat cycles without noise were selected to obtain the average RVSP.

*Rat RVSP and pulmonary artery pressure (PAP) measurement:* rats were anesthesized with alfaxalone (30mg/kg), midazolam (5mg/kg) and fentanyl (0.2mg/kg). A pre-curved catheter was inserted through the right jugular vein into the RV and into the pulmonary artery (PA). Hemodynamic data was recorded in the RV and PA using a Biopac MP36R System and Biopac's Aqcknowledge 4.1.0 software. For each rat, at least 30 seconds of continuous and stable heartbeat cycles without noise were selected to obtain the average RVSP and PAP.

*Necropsy and tissue collection:* After hemodynamic measurement, blood was extracted from the inferior vena cava and poured into EDTA tubes at RT until centrifuged for 10 minutes at 4500 rpm for plasma collection. Animals were then culled and perfused via the RV with saline solution. Next, heart and lungs were dissected and weighed. Hearts and lungs were dissected and weighed. The atria were removed, and the RV was carefully separated from the left ventricle plus septum (LV+S). RV and LV+S were weighed individually, and the RV/(LV+S) ratio was calculated as the RV hypertrophy index (Fulton's index). Organs were frozen in liquid nitrogen and stored at -80°C until protein extraction. In addition, left lung lobule was inflated with 4% paraformaldehyde (PFA) and stored in PFA with RV and left ventricle (LV) at 4°C for 24-48 hours. Then, left lung, RV and LV were washed 3 times for 5 minutes before being placed in ethanol 70% for paraffin processing.

*Lung tissue histology and immunohistochemistry:* Transverse 4 µm-thick lung sections were processed. Elastic van Gieson (EVG) staining was performed following manufacture's conditions. Peripheral vessels less than 100 µm in diameter were counted at ×40 magnification objective, and pulmonary vascular remodeling was expressed as the % of vessels with double elastic lamina (>75% of the circumference) to total vessels counted. For immunohistochemistry staining, a heat induced antigen retrieval (HIER) procedure was conducted. Incubation with the primary antibody anti-SMA (Abcam ab5695) at 1:400, then incubated with a HRP-conjugated polymer (Envision 4003, Agilent) as secondary antibody. Immunostaining was visualized with 3,3´- diaminobenzidine (DAB) and counterstained with DAKO hematoxylin (S3301). For image analysis the samples were digitalized with a scanner (AxioSan Z1 from Zeiss®).

***Echocardiographic analysis***

Echocardiography in mice and rats were performed to evaluate the pulmonary artery (PA) main trunk as well as cardiac dimensions and function. Transthoracic echocardiography was blinded performed by an expert operator at the CNIC Advanced Imaging Unit, using a high-frequency ultrasound system (Vevo 2100, Visualsonics Inc., Canada) with a 30-MHz linear probe. Two-dimensional (2D) and M-mode (MM) echography were performed at a frame rate above 230 frames/sec, and pulse wave Doppler (PW) was acquired with a pulse repetition frequency of 40 kHz. Mice were lightly anesthetized with 0.5-2% isoflurane in 100% oxygen, adjusting the isoflurane delivery trying to maintain a light anesthesia plane (heart rate 450±50 bpm). Mice were placed in supine position using a heating platform and warmed ultrasound gel was used to maintain normothermia. A base apex electrocardiogram (ECG) was continuously monitored. Images were transferred to a computer and were analyzed off-line using the Vevo 2100 Workstation software. For LV systolic function assessment, parasternal standard, long and short axis views (LAX and SAX view, respectively) were acquired. LV ejection fraction (EF), LV fractional shortening (FS), and LV chamber dimensions were calculated from these views. RV systolic function was assessed using tricuspid annular plane systolic excursion (TAPSE), obtained from an M-mode 4-

chamber apical view by measuring the maximum lateral tricuspid annular movement. RV dimensions were evaluated by measuring RV diameter. In addition, the mitral valve (MV) inflow pattern was acquired using PW Doppler echography in the 4-chamber apical view to assess diastolic function. Early and late diastolic velocity peak wave (E and A, respectively), the E/A ratio and isovolumetric relaxation time (IVRT) were measured. Cardiac Output (CO) was also measured and PVR values were calculated accordingly (mPAP = PVR x CO). To assess pulmonary pressures, the PA flow was measured from a 2D SAX at the level of the great vessels, optimized to visualize the PA crossing the aorta and parallel to the ultrasound beams. PW Doppler was displayed just at the beginning of the PA. The PA acceleration time (AT) and ejection time (ET) were measured, and the ratio was calculated.

##### *Wire myography for arterial contractility studies*

Pulmonary artery's function was assessed by wire myography. Briefly, animals were sacrificed and left lung was completely excised, washed, and preserved in Krebs-Henseleit solution (KHS: 115 mM NaCl, 2.5 mM $CaCl_2$, 4.6 mM KCl, 1.2 mM $KH_2PO_4$, 1.2 mM $MgSO_4$, 25 mM $NaHCO_3$, 11.1 mM glucose, and 0.01 mM EDTA). After that, the main pulmonary artery and secondary intrapulmonary branches were carefully dissected and cleaned of pulmonary surrounded pulmonary tissue and cut into ~2 mm long segments. Pulmonary artery rings were mounted on two tungsten 25µm diameter wires in a wire myograph system (620M, Danish Myo Technology A/S, Hinnerup, Denmark) and immersed in 37°C KHS with constant gassing (95% $O_2$ and 5% $CO_2$). Wire myography was performed as previously described.[6] Optimal vessel distension was determined by normalization using the Laplace Equation (Tension = [pressure × radius]/thickness) to calculate the position at which the tension was equivalent to an intraluminal pressure of 25 mmHg (L100);[7] vessels were then set up to the optimal tension (physiological distension, 0.9 of L100). After equilibration for 30 min, vasoconstriction was studied by exposing the pulmonary artery rings first to 80 mM KCl. Vasodilation was assessed by examining the response to increasing doses of mirabegron (from 0.1 nM to 10 µM) in segments previously contracted with phenylephrine 1 µM (Sigma-Aldrich). Drug treatments were separated by extensive washes and a stabilization period of at least 30 minutes.

##### *Human COPD-PH samples*

Human lung tissue samples were obtained from lung transplant recipients with chronic obstructive pulmonary disease (COPD) with or without PH.[8] Control samples were obtained from donors undergoing surgical lung resection for lung cancer as part of a study of histology in smokers.[9] Studies were approved by the Ethics Committee of Hospital Clinic, Barcelona, Spain, and all patients provided written informed consent. Lung tissue samples were collected prospectively and processed after surgery. Briefly, immediately after resection, fresh 3-4 mm thick lung tissue slices were washed 3 times (10 minutes each) in cold PBS in order to distend collapsed areas of parenchyma and clean off embedded blood. Paraffin-embedded lung tissue blocks were cut into 5 µm sections and processed for immunohistochemistry.

***Cell culture***

Human pulmonary arterial endothelial cells (HPAECs; Promocell: C-12241) and human pulmonary arterial smooth muscle cells (HPASMCs; Promocell: C-12521) were cultured with Endothelial Cell Growth Medium (PromoCell: C-22010) and Smooth Muscle Cell Growth Medium 2 (Promocell: C-22162), respectively. Cells were used between passages four and seven and cultured in a normal pressure incubator, either in normoxia (21% $O_2$, 5% $CO_2$, 37ºC) or hypoxia (2% $O_2$, 5% $CO_2$, 37ºC) for the timepoints indicated in the manuscript. All experiments were repeated at least 3 times. Cells were incubated either with mirabegron (MB, 0.1µM, Santa Cruz Biotechnologies: sc-211912), as an β3-AR agonist; L-Nitro-Arginine Methyl Ester (L-NAME, 100uM, Selleckchem: S2877) as an inhibitor of nitric oxide synthase and /or Genipin (10µM; Sigma: G4796) as an inhibitor of UCP2.

***Tissue and cells immunofluorescence***

*Tissue immunofluorescence:* lungs and hearts were embedded in paraffin and cut at 5 µm in the microtome (Leica). Paraffin was melted in the incubator at 60 °C for 20 minutes. After that, slides were incubated with xylol for 20 minutes at room temperature (RT). Slides were hydrated with alcohols following this sequence: Ethanol 100% (5 minutes), ethanol 96% (5 minutes), ethanol 70% (5 minutes) and $H_2O$d (5 minutes). For antigen unmasking, slides were submerged in citrate buffer 10mM pH6 (sodium citrate dihydrate 4mM and citric acid dihydrate 6 mM) for 30 minutes in the microwave. After that, slides were cooled in citrate buffer for 30 minutes at RT. For membrane permeabilization, slides were washed with PBS for 5 minutes, incubated with PBS-Triton 0,3% for 7 minutes and washed again 3 times with PBS for 5 minutes. For blocking, cuts were incubated for 1 hour in blocking buffer (Goat Serum 5% in PBS) at RT. Slides were incubated overnight at 4 °C in wet chamber with primary antibodies: mouse anti-SMA (Abcam, ab7817; 1:200), rabbit anti-CD31 (Abcam, ab28364; 1:200), rat anti-CD31 (DIANOVA, DIA-310; 1:50), rabbit anti-Ki67 (Thermofisher, PA519462; 1:250), rat Ki67-eFluor660 (eBioscience, #50-5698-82; 1:100), rabbit anti-vWF (Dako, A0082, 1:100), rabbit anti-β3-AR· (Genetex, GTX108200, 1:100), rabbit anti-Glut1 (Millipore, #07-1401, 1:100), mouse anti-SMA-Cy3 (Sigma, C6198, 1:400), mouse anti-SMA-HRP (Sigma, SAB4200679, 1:1000), rabbit anti-SMA (Abcam, ab150301, 1:200), diluted in antibody buffer (Tween20 0,05%; Goat Serum 2% in PBS). The following day, slides were washed 3 times for 5 min with PBS at RT in agitation. Slides were incubated with proper secondary antibody conjugated with a fluorochrome for 1 hour at RT in obscurity: goat anti-rat-Cy3 (Jackson ImmunoResearch, 112-166-062, 1:500), goat anti-mouse-Alexa488 (Thermofisher, A-11029, 1:500), goat anti-rabbit-Alexa546 (Thermofisher, A-11035, 1:500), goat anti-rabbit-Alexa647 (Thermofisher, A-21443, 1:500), anti-rabbit-HRP (Agilent, K4003, 1:500). Next, slides were washed 3 times for 5 minutes with PBS. After that, slides were incubated with DAPI (1:10000 in PBS) for 5 minutes at RT, in agitation and obscurity. Slides were mounted using Fluoroshield (4958-02, Thermofisher Scientific) mounting medium and cover slips of 1 mm. Peripheral vessels less than 30μm (mouse lungs) or 100μm (rat lungs) were considered for quantifications.

*Cell immunofluorescence*: human pulmonary artery smooth muscle cells (HPASMCs) cultured on round cover glass in 24-well and maintained under hypoxic conditions in a hypoxic chamber (2% $O_2$) for 24 hours. Control cells were kept in normoxia conditions. Half of wells were treated with mirabegron in concentration $10^{-7}$M and half was kept without treatment as a control. Then, cells were fixed with 4% paraformaldehydes in PBS for 15 minutes. Cells were washed 2 times with PBS. For membrane permeabilization cells were incubated with Triton 0,5% in PBS for 10 minutes at RT. Then, cells were blocked with blocking solution (goat serum 5% in Tris Buffer Solution-Tween (TBST) for 30 minutes at RT. Samples were incubated overnight at 4˚C with primary antibody Anti-Ki67 (1:200; ab16667, Abcam). After 3 washed with PBS, cells were incubated with secondary antibody conjugated with a fluorochrome anti-rabbit-Alexa-546 (Thermofisher, A-11035, 1:500) for 1 hour at RT. Cells were washed 3 times with PBS and incubated for 5 minutes with DAPI (1:10000 in PBS) or Hoechst (1:10000 in PBS) and washed again 3 times with PBS. Stained cells were mounted in Fluoromount G imaging medium (4958-02, Thermofisher Scientific). In addition, to study mitochondria and oxidative stress in cells we added *in vitro* MitoTracker Red CMXRos (M7512, Thermofisher Scientific; 200 nM) and CellRox Green Reagent (C10444, ThermoFisher Scientific; 5 μM) to each well for 30 minutes at 37˚C. Next, cells were washed 3 times with PBS and fixed with PFA 4% for 10 minutes at RT. After fixation, the cells were rinsed in PBS several times. Cells were permeabilized with 0,1% Triton in PBS 1x for 3 minutes and then rinsed twice with PBS. Then, cells were incubated for 10 minutes with DAPI (1:100 in PBS) and washed again 3 times with PBS. Finally, Stained cells were mounted in Fluoromount G imaging medium (4958-02, Thermofisher Scientific). Additionally, levels of cellular ROS were quantified using CellROX Green (ThermoFisher Scientific: C100448) and used according to manufacturer's instructions. Briefly, cells were incubated with 5µM of the probe at 37˚C, for 30 minutes. Cells were washed 3 times with PBS prior to fixation and nuclear staining with DAPI, and the visualized using the Zeiss700 laser scanning confocal microscope. Confocal imaging was performed on a Leica SP5 confocal microscopy and Leica SP8 STED confocal microscopy. The appropriate lasers were used to see DAPI, Hoechst-33342, 488, 520, 546, 590 and 647 nm excitations. x63 objective with immersion oil was used. Images were analysed using Fiji ImageJ software.

*HPASMC proliferation assays:* to assess HPASMC proliferation, cells were first starved o/n, then serum-free/factor-free medium was replaced by HPASMC complete medium with or without agonists/inhibitors, or alternatively, by conditioned medium of HPAECs treated or not with agonists/inhibitors and exposed to hypoxia. 20 hours after exposure to hypoxia, EdU (5-ethynyl-2'-desoxyuridine; ThermoFisher Scientific: C10340) was added to HPASMCs at a final concentration of 5µM). Cells were left for an additional 4 hours in hypoxia before washing and fixation. Cells were then immunostained for Ki67 (ThermoFisher Scientific: PA5-19462) and EdU detection was carried out according to manufacturer's instructions.

***Cell-based mitochondrial respiration experiments***

Oxygen-consumption rates (OCR) were measured at 37°C using a Seahorse XF96 analyser. Briefly, $2.0\times10^4$ HPAECs/well were seeded in an XFE-24 Seahorse plate (Seahorse Biosciences, 102340) precoated with 0.2% gelatine and allowed to adhere for 24hrs at 37°C with 5% $CO_2$. Medium was then renewed with or without agonists/inhibitors and cells were placed in hypoxia (2% $O_2$, 5% $CO_2$, 37°C) for 24 hours. Unbuffered DMEM without bicarbonate (Sigma, D5030) supplemented with 15 mM glucose, 2 mM sodium pyruvate and 1 mM glutamine – hereon named Seahorse medium - was also placed overnight in the hypoxia chamber to equilibrate. Medium was removed from cells, and cells were washed with Seahorse medium (with pH adjusted to 7.4 just prior to use) and then incubated with 175µL of seahorse medium containing or not the corresponding agonists/inhibitors in a 37°C non-$CO_2$

incubator for 30 minutes. Oligomycin (Sigma: O4876), FCCP (Sigma: C2920), and rotenone + antimycin A (Sigma: R8875 and SigmaA8674) were all used at a final concentration of 1µM. OCR measurements were then normalized to cell content, which was assessed by fluorometry using ReadyProbes™ Cell Viability Imaging Kit, Blue/Red (R37610).

***Western blot***

Tissue and cells samples were homogenized with RIPA buffer 10X (Merck, REF: 20-188) supplemented with phosphatase inhibitor (4X) and protease inhibitor (4X) (cOmplete™ Protease Inhibitor Cocktail and PhosSTOP™, Roche, respectively) using Qiagen Tissue Lyser LT tubes holder that was at -80˚C. Each sample were placed in a 2 mL tube (Sarstedt Ref. 72693465) with 2 steel beads (Qiagen Ref. 69989 5mm). First, 15 minutes at 20 oscillations per 1/s was done. At the end of this time, the tissue was almost completely lysed. Tissue lysate was centrifuged 30 minutes, 16000g, 4˚C. Finally, supernatant was collected for protein quantification. Protein concentration was determined by BCA Protein Assay Kit from Thermo Scientific™ (Ref. 23225) using bovine serum albumin (BSA) as the standard. Protein samples derived from mouse tissues (30–40 µg) or cultured cells (20–30 µg) were heated at 95°C for 5 minutes and loaded in 7.5 – 15% acrylamide gel for their separation in SDS-PAGE (Tris-glycine gels with Tris/glycine/SDS buffer) and transferred onto nitrocellulose membranes using 0.2 µm Turbo™ Transfer System (BioRad, Ref. 1704158) following manufacturer's protocols. Blots were stained with Ponceau Red Solution for 3-4 minutes, washed with TBS-Tween (0,2%) 1X (TBST 1X) and incubated in blocking buffer (5% BSA in TBST) at RT for 1 hour. Primary antibody incubation was performed overnight at 4ºC using: mouse anti-eNOS (1:1000; lab 610296, BD Transd.), rabbit anti-UCP2 (1:2000; GTX132072, Genetex), rat anti-CD31 (1:1000; DIA-310, Dianova), anti-Tom20 (1:100; Thermofisher PA5-52843) and mouse anti-VINCULIN (1:2000; V4505, Sigma-Aldrich). Afterwards, membranes were washed with TBST 1X and incubated for 1 hour at RT with the corresponding secondary antibody HRP-conjugated, either Anti-rabbit (1:5000; P0448, Dako), Anti-mouse (1:5000; P0447, Dako) or Anti-rat (1:5000; 3010-05, SouthernBiotech) diluted in BSA 1% TBST. Membranes were incubated with Immobilon® Western Chemiluminescent HRP Substrate (MERCK©) and visualized with an ImageQuant LAS 4000 mini–Biomolecular Imager (GE-HealthCare©). Optical densities of individual bands were measured using Fiji ImageJ software and protein expressions were normalized by vinculin protein expression.

***Flow cytometry***

Right lung lobes were collated into cold saline solution, then were placed on petri dish top, and minced using scissors and scalpel. Minced lungs were digested with digestion buffer (1mg/mL Collagenase II (Worthington, LS004176), 2mg/mL Dnase (Roche, 11284932001), 5mM $CaCl_2$ in PBS). Homogenate's lungs were incubated in a bacterial shaker at 37°C, 100 r.p.m., for 30 minutes. Then, Homogenates were passed through a 40µm cell strainer, washed with FBS 5%, PBS and centrifugated at 500g for 6 minutes. After that, RBC lysis Buffer (Invitrogen, 00433357) was added and incubated for 3 minutes at RT. Additionally, FBS 5%, PBS were added and they were centrifuged for 6 minutes at 500g. Pellets were resuspended in FBS 5%, PBS with anti-CD31-APC (BD Pharmingen, 551262) and Anti-CD45-V450 (BD Biosciences, 560501) and incubated for 20 minutes in ice. Then, they were centrifuged for 6 minutes at 500g and pellets were resuspended in FACS buffer (FBS 5%, PBS) to take them to the cytometer. Only $CD31^+$ and $CD45^-$ cells were sorted. At the end of the protocol, usually 200,000 / 300,000 ECs were obtained in lung right lobes.

***RNA extraction and qPCR***

FACs sorted cells were pelleted and lysed in buffer RLT (RNAeasy Micro kit – Qiagen, 74004) and RNA extracted according to the manufacturer instructions and stored at -80°C. Subsequently, the total RNA obtained from each sample was reverse-transcribed into cDNA using the High-Capacity cDNA Reverse Transcription Kit with RNase Inhibitor (Thermo Fisher, 4368814). cDNA was used for qRT-PCR with Sybr Green Master Mix and gene specific primers (10 µM) (Supp. Table 1) in an AB7900 real-time thermocycler (Applied Biosystems). Hprt (*Hypoxanthine Phosphoribosyltransferase 1*) expression was used as a housekeeping gene. Finally, gene expression was calculated using the $\Delta\Delta$Ct method and the ribosomal 36b4 gene used as reference gene for data normalization.

***ELISA (Enzyme Linked ImmunoSorbent Assay)***

cAMP and cGMP levels in HPAECs and HPASMCs were determined using the cAMP and cGMP Elisa Kits from Cayman Chemicals (581001 and 581021, respectively). In addition, ELISA kits were used to measure nitrite plasma levels (Invitrogen cat. EMSNO) and endothelin (ET-1) (R&D DET100) following the protocol from the manufacturer in mice.

##### *Statistical analyses*

Experimental data are presented as mean ± standard error of the mean (SEM) and were analysed with Prism software (GraphPad Prism) and Fiji ImageJ software version 1.54i. Comparisons between two groups were made by unpaired two-tailed Student t-test and paired two-tailed Student t-test. Comparisons between more than two groups were made by two-way ANOVA followed by multiple comparisons test as described on each figure legend depending on the experimental setting. Statistical significance was defined as *p<0.05, **p<0.01, ***p<0.005, ****p<0.001, #p<0.05, ##p<0.01, ###p<0.005, ####p<0.0001.

#### SUPPLEMENTAL TABLES

**Supp. Table 1. Primers for AdrB3 recognize both mouse and human sequences and fall within the deleted region of the mouse AdrB3 KO.** In order to compare directly expression levels of AdrB3 in the mouse KO and the OE of the human AdrB3.

| Gene | Fwd primer | Rev Primer |
|---|---|---|
| AdrB3 | CCCATCATGAGCCAGTGGT | GGGGAAGGTAGAAGGAGACG |
| 36b4 | ACTGGTCTAGGACCCGAGAAG | TCCCACCTTGTCTCCAGTCT |

**Supp. Table 2. Parameters determined by echocardiography in MCT-induced PH in rats and subjected to different treatment schemes with mirabegron.** HR: Heart rate; LVEF: Left ventricle ejection fraction; PAAT: Pulmonary artery acceleration time; PA. Diam.: pulmonary arterial diameter; PVR: Pulmonary vascular resistance; PV-VTI: pulmonary valve velocity time integral; RVCO: RV cardiac output; RV Diam.: RV diameter; RVEF: RV ejection fraction; RVOT: RV outflow tract; RV Stroke Vol.: RV stroke volume; TAPSE: Tricuspid annular plane systolic excursion.

| | Units | Mean ± SEM | | | | Adjusted p-value in relation to control | | |
|---|---|---|---|---|---|---|---|---|
| | | **Control** | **MCT** | **MCT+Treat.** | **MCT+Prev** | **MCT** | **MCT+Treat** | **MCT+Prev.** |
| PV-VTI | mm | 44.96±2.66 | 25.85±2.45 | 23.87±2.40 | 32.22±3.67 | <0.0001 | <0.0001 | 0.008 |
| PAAT | ms | 32.23±1.19 | 19.09±2.03 | 17.82±1.03 | 17.99±1.02 | 0.0001 | 0.0001 | 0.0001 |
| TAPSE | mm | 2.412±0.15 | 1.86±0.21 | 2.186±0.17 | 2.35±0.30 | 0.278 | 0.717 | 0.833 |
| RVOT | mL/min | 3.751±0.13 | 3.77±0.10 | 3.720±0.15 | 3.92±0.08 | 0.999 | 0.995 | 0.594 |
| PA. Diam. | mm | 2.475±0.10 | 2.37±0.20 | 2.731±0.13 | 2.77±0.14 | 0.935 | 0.470 | 0.350 |
| RV Diam. | mm | 3.965±0.20 | 5.50±0.24 | 5.794±0.32 | 5.89±0.42 | 0.0044 | 0.0005 | 0.0002 |
| RV wall | mm | 0.793±0.03 | 1.35±0.11 | 1.478±0.11 | 1.27±0.10 | 0.0006 | <0.0001 | 0.0020 |
| RVCO | mL/min | 56.16±10.56 | 27.53±4.50 | 44.8±6.99 | 41.81±9.32 | 0.069 | 0.639 | 0.506 |
| RVEF | % | 56.42±2.28 | 40.74±3.40 | 45.16±3.26 | 44.68±4.35 | 0.008 | 0.067 | 0.048 |
| RV Stroke Vol | mL | 135.90±17.94 | 97.22±14.43 | 118.80±16.30 | 102.00±13.60 | 0.223 | 0.782 | 0.288 |
| PVR | mmHg/mL/min | 0.19±0.02 | 0.47±0.06 | 0.311±0.064 | 0.240±0.03 | 0.0068 | 0.1704 | 0.7701 |
| HR | bpm | 341.0±4.20 | 344.4±9.40 | 335.7±10. | 338.6±9.50 | 0.988 | 0.947 | 0.996 |
| LVEF | % | 65.99±2.20 | 64.44±3.90 | 64.69±5.00 | 65.52±2.93 | 0.980 | 0.988 | 0.999 |

#### SUPPLEMENTAL FIGURE LEGENDS

**Supplementary Figure 1. Mice lacking β3-AR present a more severe PH phenotype.**
**(A)** Ratio between left lung lobe and body weight of normoxic and hypoxic WT and AdrB3$^{KO}$ mice. **(B)** Ratio between RV and body weight (BW) of normoxic and hypoxic WT and AdrB3$^{KO}$ mice. **(C)** Ratio between left ventricle (LV) + septum (S) and body weight of normoxic and hypoxic WT and AdrB3$^{KO}$ mice. **(D)** Quantification of CD31 protein levels in lungs and representative image of western blot. **(E)** Gating of dissociated lungs of normoxic and hypoxic WT and AdrB3$^{KO}$ mice to identify EC population (CD31+, CD45-). **(F)** Chart representing lung EC frequency determined by FACS analysis. All data are represented as mean ± s.e.m. Each dot in charts represents an individual mouse. Two-way ANOVA with Sidak's multiple comparisons test. Asterisks indicate p-values between normoxic and hypoxic mice of the same genotype. *p<0.05, **p<0.01, ****p<0.001.

**Supplementary Figure 2. Recovery of β3-AR in the endothelium improves features of hypoxia-induced PH in KO mice. (A)** Gating of dissociated lungs of normoxic AdrB3$^{KO}$ and AdrB3$^{EC}$ mice to identify the frequency of transgene expression (R26$^{-LSL-hB3-AR-IRES-GFO}$) in ECs (CD31$^{+}$, CD45$^{-}$) and hematopoietic cells (HC; CD45$^{+}$, CD31$^{-}$). Quantifications are presented in Fig. 2C. **(B)** Ratio between left lung lobe and body weight of normoxic and hypoxic AdrB3$^{KO}$ and AdrB3$^{EC}$ mice. **(C)** Ratio between RV and body weight of normoxic and hypoxic AdrB3$^{KO}$ and AdrB3$^{EC}$ mice. **(D)** Ratio between left ventricle + septum and body weight of normoxic and hypoxic AdrB3$^{KO}$ and AdrB3$^{EC}$ mice. All data are represented as mean ± s.e.m. Each dot in charts represents an individual mouse. Two-way ANOVA with Sidak's multiple comparisons test. Asterisks indicate p-values between normoxic and hypoxic mice of the same genotype. **p<0.01, ***p<0.005, ****p<0.001.

**Supplementary Figure 3. Restoration of β3-AR in SMC had no effects on hemodynamic measures but a downward trend in vessel remodeling. (A)** Hypoxia-induced pulmonary hypertension experimental setup. **(B, C)** Representative recording profiles and quantification of RVSP. **(D)** Fulton's index reflecting RV hypertrophy. **(E)** Ratio between left lung lobe and body weight of normoxic and hypoxic AdrB3$^{KO}$ and AdrB3$^{SMC}$ mice. **(F)** Ratio between RV and body weight of normoxic and hypoxic AdrB3$^{KO}$ and AdrB3$^{SMC}$ mice. **(G)** Ratio between left ventricle + septum and body weight of normoxic and hypoxic AdrB3$^{KO}$ and AdrB3$^{SMC}$ mice. **(H)** Representative images of mouse lung sections with immunostained for SMA. Arrowheads indicate fully muscularized arterioles with less than 30µm diameter. Inserts depict representative arterioles. **(I)** Quantification of density of fully muscularized arterioles in lung sections. **(J)** Quantification of proliferating lung SMCs. **(K)** Representative confocal images of Ki67 (red) and SMA (green) immunostaining in mouse lung sections. Asterisks indicate proliferative $Ki67^{+}$ SMCs. All data are represented as mean ± s.e.m. Each dot in charts represents an individual patient/mouse. Two-way ANOVA with Sidak's multiple comparisons test. Asterisks indicate p-values between normoxic and hypoxic mice of the same genotype and hashtags indicate p-values between genotypes. *p<0.05, **p<0.01, ***p<0.005, ****p<0.001, #p <0.05.

**Supplementary Figure 4. Mirabegron improves cardiac structure and function in hypoxic mice. (A)** Heat map representing change in percentage of several parameters determined by echocardiography of hypoxia-exposed mice treated with DMSO, MB2 or MB10 in drinking water and parameters determined by echocardiography in hypoxia-exposed mice and subjected to treatment with mirabegron. PAAT: Pulmonary artery acceleration time; PA. Diam.: pulmonary arterial diameter; PV-VTI: pulmonary valve velocity time integral; RV Diam.: RV diameter; RVOT: RV outflow tract; TAPSE: Tricuspid annular plane systolic excursion. **(B-G)** Charts with the quantification of the parameters indicated in (A). **(H)** Quantification of the proportion of RV area positive for Glut1 immunostaining. **(I)** Representative confocal images of mouse RV sections immunostained for Glut1 (red) and WGA (Wheat Germ Agglutinin; green). All data are represented as mean ± s.e.m. Each dot in charts represents an individual patient/mouse. One-way ANOVA with Dunnett's multiple test comparisons. Asterisks indicate p-values between normoxic and hypoxic mice. *p<0.05, ***p<0.005. PA – pulmonary artery; PAAT - pulmonary artery acceleration time; PV-VTI - pulmonary venous velocity time integral; RVOT – RV outflow tract; TAPSE - tricuspid annular plane systolic excursion.

8. Peinado VI, Gómez FP, Barberà JA, Roman A, Angels Montero M, Ramírez J, Roca J, Rodriguez-Roisin R. Pulmonary vascular abnormalities in chronic

obstructive pulmonary disease undergoing lung transplant. *The Journal of Heart and Lung Transplantation*. 2013;32:1262–1269.

9. García-Lucio J, Argemi G, Tura-Ceide O, Diez M, Paul T, Bonjoch C, Coll-Bonfill N, Blanco I, Barberà JA, Musri MM, et al. Gene expression profile of angiogenic factors in pulmonary arteries in COPD: relationship with vascular remodeling. *American Journal of Physiology-Lung Cellular and Molecular Physiology*. 2016;310:L583–L592.

**Supplementary Figure 1**

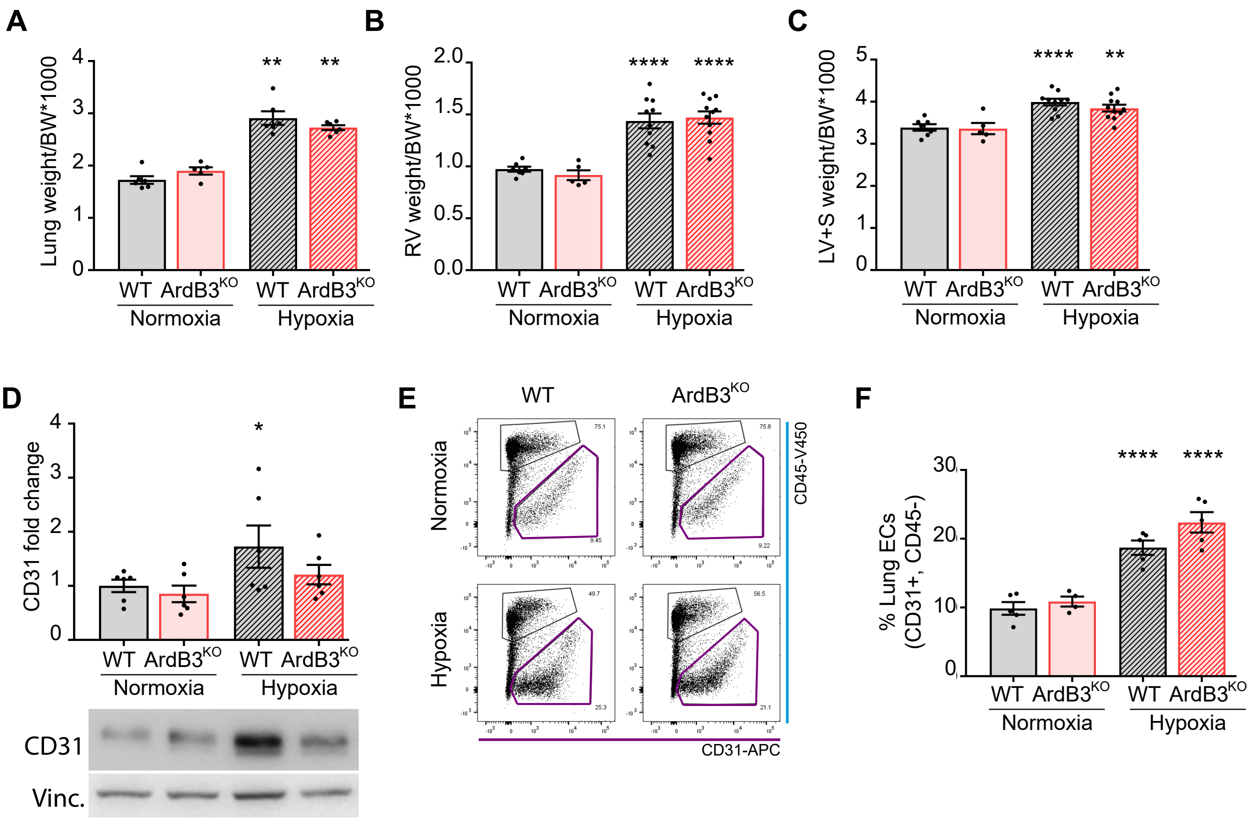

### Supplementary Figure 2

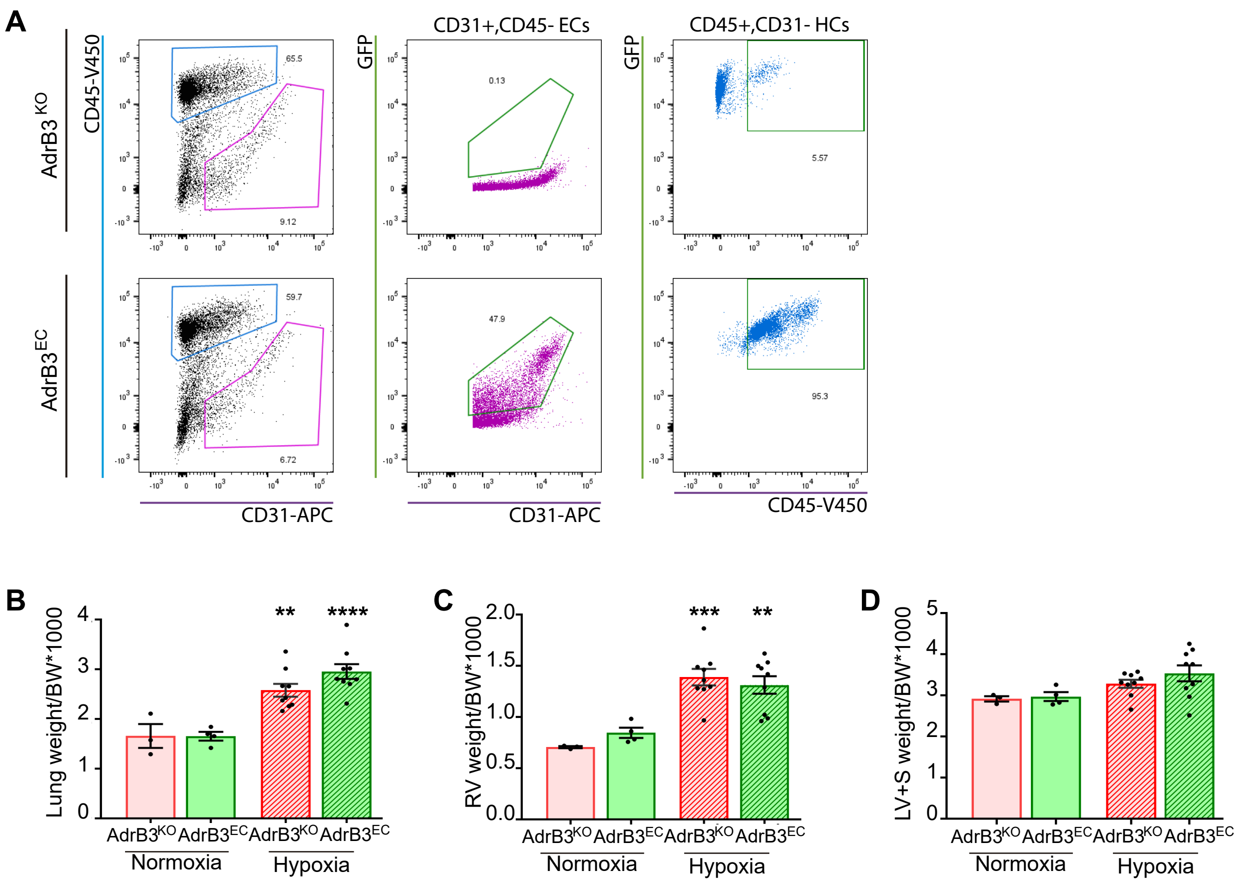

#### Supplementary Figure 3

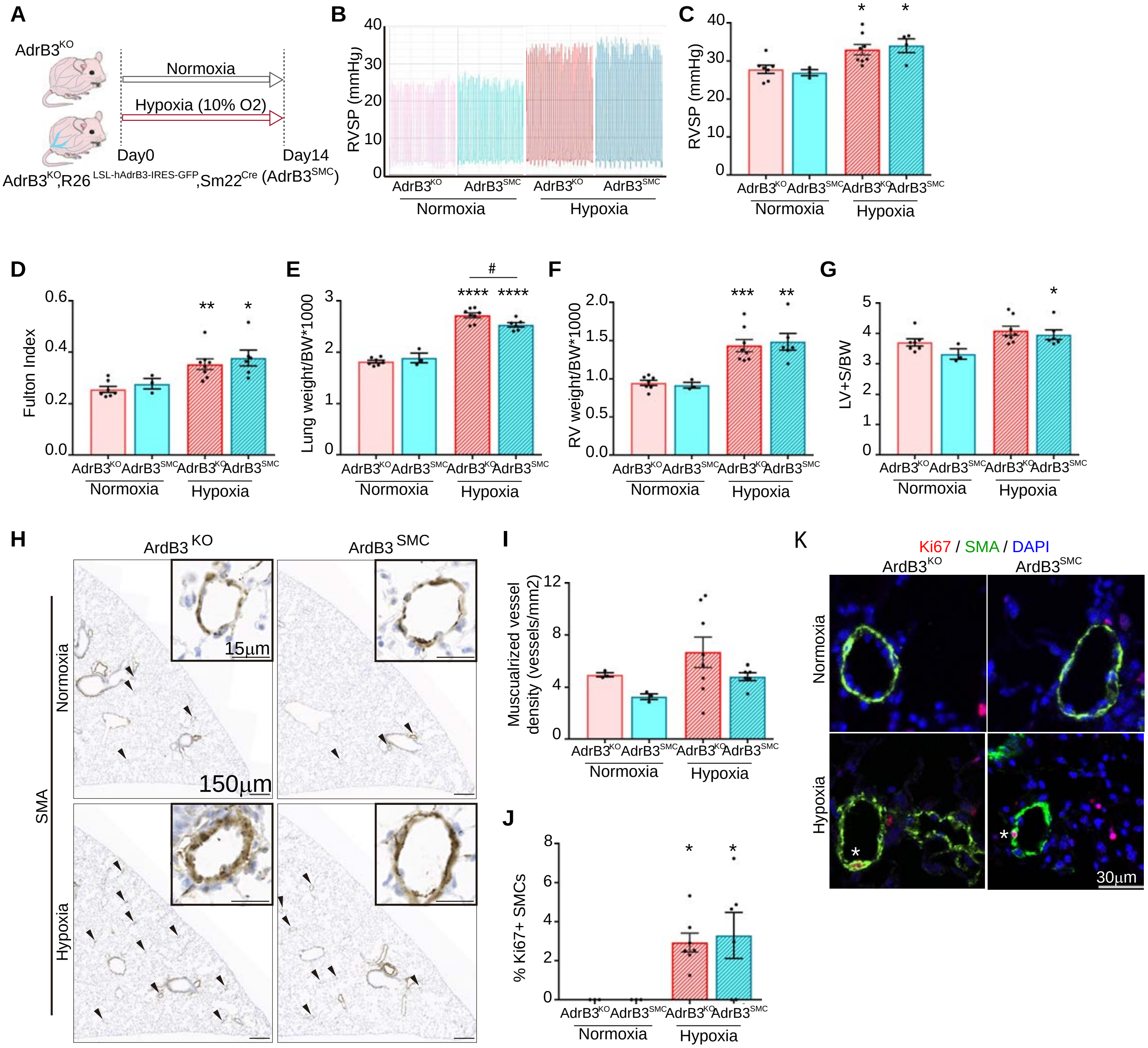

### Supplementary Figure 4

A

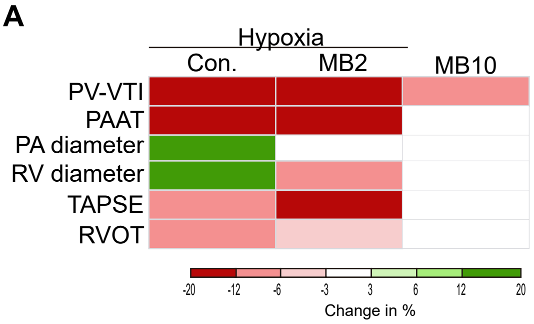


| | Units | Mean ± SEM | | | | Adjusted p-value in relation to normoxia | | |
|---|---|---|---|---|---|---|---|---|
| | | **Normoxia** | **DMSO** | **MB2** | **MB10** | **DMSO** | **MB2** | **MB10** |
| PV-VTI | mm | 23.95±1.570 | 19.60±1.050 | 20.88±1.420 | 21.24±1.030 | 0.0008*** | 0.029* | 0.011* |
| PAAT | ms | 18.92±1.580 | 16.25±0.910 | 16.39±0.650 | 18.36±0.720 | 0.213 | 0.406 | 0.970 |
| TAPSE | mm | 0.778±0.048 | 0.711±0.066 | 0.639±0.085 | 0.775±0.062 | 0.779 | 0.412 | 0.999 |
| RVOT | mm | 1.606±0.074 | 1.723±0.054 | 1.667±0.098 | 1.549±0.104 | 0.534 | 0.927 | 0.927 |
| PA Diam. | mm | 0.839±0.049 | 0.952±0.042 | 0.840±0.071 | 0.857±0.041 | 0.205 | 0.999 | 0.985 |
| RV Diam | mm | 1.595±0.077 | 1.848±0.121 | 1.459±0.099 | 1.593±0.138 | 0.272 | 0.826 | 0.999 |

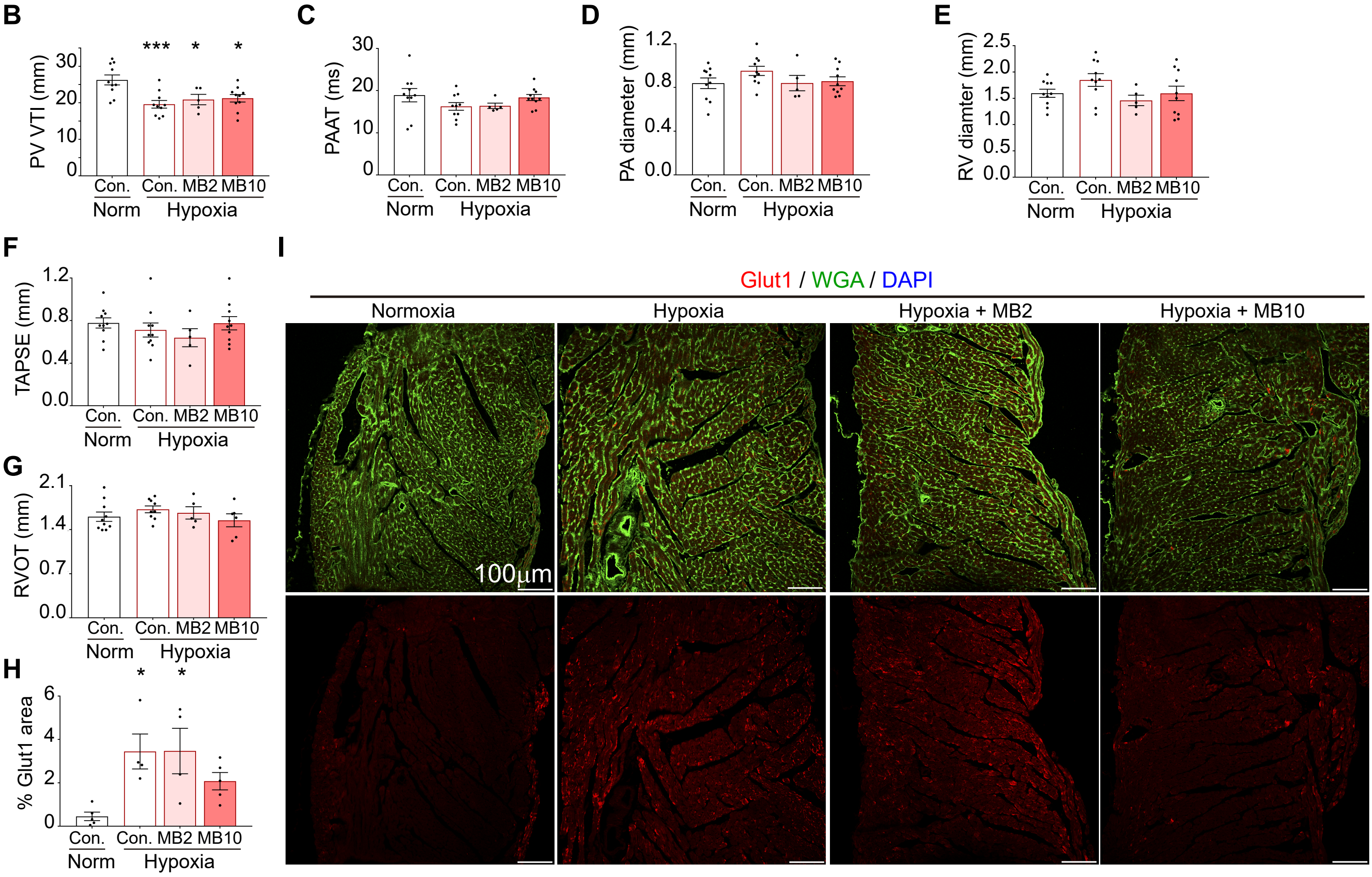
